# Rsm-mediated post-translational control of the *Pseudomonas putida* Type VI Secretion System

**DOI:** 10.64898/2026.07.10.737732

**Authors:** Cristina Civantos, Carmen Paredes, Marina Murillo-Torres, João Botelho, María Antonia Sánchez-Romero, Luke P. Allsopp, Patricia Bernal

**Author notes:** **Author’s Note:** Four supplementary figures and four supplementary tables are attached.

## Abstract

The Type VI secretion system (T6SS) is a bacterial nanoweapon that injects toxic effectors into prokaryotic and eukaryotic cells. It is widely found among gram-negative bacteria and provides a significant fitness advantage in interbacterial competition. *Pseudomonas putida* KT2440 possesses three T6SS clusters (K1-, K2- and K3-T6SS) that combat phytopathogens. This makes this strain a potent biocontrol agent that protects plants from pathogens and can be further enhanced by a better understanding of its T6SS regulation. Although the core components of T6SS are conserved, the elements controlling its regulation differ among bacterial species. T6SS activity is regulated by various factors acting at different levels, from transcription to post-translational modification, to ensure precise control of its activity. Here, we demonstrate the critical importance that the three Rsm proteins, RsmIEA, have in controlling the K1-T6SS structural components and related orphan elements at the post-transcriptional level in *Pseudomonas putida*. We identified multiple Rsm-binding sites responsible for directly repressing the translation of T6SS proteins (Hcp1 and Hcp5) and their associated effectors (Tke2 and Tke7). Derepression of K1-T6SS mRNA in the *rsmIEA* mutant led to enhanced translation and expression of the K1-T6SS components and effectors, and critically increased the number of cells in the population with assembled T6SS. This results in a greater capacity to secrete toxins and kill prey cells via the T6SS-dependent mechanism. Finally, we demonstrate the K1-T6SS ability to kill environmental pathogens, including *Salmonella enterica* and *Erwinia amylovora*.

## INTRODUCTION

In microbial communities, individuals of different species use strategies to eliminate each other, acting as competitors to control their ecological niches. The Type VI Secretion System (T6SS) is a protein nanomachine that functions as a poisoned harpoon, allowing bacteria to outcompete bystanders and foes. T6SS-producing bacteria inject toxic effector proteins into prey cells in a contact-dependent manner whilst simultaneously co-expressing cognate immunity proteins to neutralise the toxicity of sister cell effectors and prevent self-intoxication (Wohlfarth *et al*., 2025). This complex nanomachine consists of 13 structural core proteins that shape the three main complexes: the membrane complex, the baseplate and the tail (Allsopp and Bernal, 2023). The tail is responsible for transporting and delivering toxic effectors and is formed by an internal tube, a spike that sits on top and a contractile sheath that surrounds this structure in the cytosol. The internal tube is made of stacked hexameric rings of Hcp proteins, while the arrow-shaped spike is formed by a trimer of VgrG proteins with a PAAR protein at its pinnacle. Upon sheath contraction, the Hcp-VgrG-PAAR-effectors complex is propelled outside the producing bacteria and inside the neighbouring prey cells (Allsopp and Bernal, 2023; Wohlfarth *et al*., 2025).

The T6SS is found in pathogenic, symbiotic, and commensal microorganisms. However, it is especially relevant in highly dense microbial communities, such as the gut and the rhizosphere, where positive and negative interactions are abundant. The T6SS is important in plant-microbe interactions (Bernal *et al*., 2018; Durán *et al*., 2021; Reyes[Pérez *et al*., 2025; Vázquez-Arias *et al*., 2025). In this work, we focus on *Pseudomonas putida*, a plant growth-promoting rhizobacterium (PGPR) well known for its biocontrol ability (Weller, 2007). In *P. putida,* the T6SS is instrumental in the elimination of severe plant pathogens, and it is important for crop plant colonisation, shaping their microbiota (Bernal *et al*., 2017, 2021; Vázquez-Arias *et al*., 2025). *P. putida* KT2440 possesses three T6SSs known as K1-, K2- and K3-T6SS; five orphan gene clusters encoding Hcp (Hcp4, Hcp5 and Hcp6); two VgrG (VgrG4 and VgrG5) proteins and five associated effectors (Tke6, Tke7, Tke8, Tke9 and Tke10) (Bernal *et al*., 2017).

The K1-T6SS cluster is constitutively active in laboratory conditions, and its expression increases in the stationary phase of growth. The system is indirectly regulated at the transcriptional level by different regulators, including RpoS, RpoN, FleQ and RetS (Bernal *et al*., 2023). This tight regulation is prevalent among bacteria that harbour T6SSs, allowing them to express and assemble this probably energetically expensive structure exclusively when required, avoiding retaliation (Hespanhol *et al*., 2023). T6SS expression has been reported to be stimulated in response to factors such as high cell density, iron depletion, acidic pH, and the presence of specific host molecules (Bernard *et al*., 2011; Sana *et al*., 2012; Wu *et al*., 2012; Storey *et al*., 2020). These signals are integrated through different global regulators and pathways at transcriptional, posttranscriptional and posttranslational levels (Hespanhol *et al*., 2023).

However, there is a limited understanding of the posttranscriptional and posttranslational regulation of the K1-T6SS in *P. putida*. In contrast, in *P. aeruginosa*, the RsmA protein (repressor of <u>s</u>econdary <u>m</u>etabolism) is a key player in suppressing the translation of T6SS transcripts (Allsopp *et al*., 2017). *P. aeruginosa* RsmA is a member of the widespread RsmA/CsrA family of RNA-binding proteins. These proteins, upon binding to mRNA, can influence translation, mRNA stability, and transcript termination in both positive and negative manners (Vakulskas *et al*., 2015). Typically, members of this family bind to transcripts through their 5’ untranslated regions or the first codons of the coding regions at specific sites known as ‘Rsm binding sites’ (Vakulskas *et al*., 2015). RsmA/CsrA proteins have a strong preference for the consensus sequence 5’-RUACARGGAUGU-3’ located in the loops of short RNA hairpins (Dubey *et al*., 2005; Duss *et al*., 2014). By binding specifically to this sequence, which closely matches the ideal 5’-AAGGAGGU-3’ Shine-Dalgarno sequence (Shine and Dalgarno, 1974; Yusupova *et al*., 2001; Ma *et al*., 2002), Rsm proteins regulate the expression of numerous genes at the level of translation (Schubert *et al*., 2007). The activity of Rsm proteins is negatively regulated by the GacS/GacA two-component system (Vakulskas *et al*., 2015). This mechanism is activated under specific, yet mostly unknown, signals that promote the transcription of small noncoding RNA molecules, such as *rsmX*, *rsmY*, and *rsmZ*. These small RNAs are characterised by the presence of stem-loops with a GGA motif in their distal region, such as the Rsm binding sites, and thus, these motifs strongly bind Rsm proteins. Therefore, sRNAs sequester Rsm proteins, preventing their action on mRNA targets (Kay *et al*., 2005).

*P. putida* contains three members of the CsrA/RsmA family known as RsmI, RsmE and RsmA. RsmA and RsmE share 54% identical residues, while RsmI shows 43% identity with the other two proteins (Huertas-Rosales *et al*., 2016). Rsm proteins play a significant role in regulating biofilm formation by repressing the translation of components involved in the synthesis of different exopolysaccharides, such as Alg, Peb, Bcs, and Pea, as well as CfcR diguanylate cyclase. Bacterial attachment in *P. putida* is predominantly modulated by RsmE. Conversely, RsmI exerts a weaker effect that is contingent upon functional RsmA, indicating a coordinated regulatory interplay among these Rsm proteins. Their effect can be counteracted by three noncoding sRNAs, *rsmX*, *rsmY* and *rsmZ,* present in this strain (Huertas-Rosales *et al*., 2016, 2017). Here we investigated the post-transcriptional regulation of *P. putida* T6SSs by the Rsm system and shed light on the complex regulatory process that allows bacteria to efficiently kill their competitors. Understanding this activity will allow us to optimise the potent biocontrol apparatus of *P. putida* for future applications in agrotechnology to boost sustainable agriculture.

## MATERIAL AND METHODS

### Bacterial strains and growth conditions

The bacterial strains used are listed in Table S1. Unless otherwise stated, all chemicals and reagents, including antibiotics, were purchased from Sigma-Aldrich. All strains were grown in Lysogeny broth (LB) (Lennox LB 5 g/L NaCl) and agar (1.5% w/v) (Sambrook et al., 1989) for routine growth with shaking at 180 RPM, as appropriate. *E. coli* and *Salmonella* strains were incubated at 37°C, phytopathogens at 28°C and *P. putida* strains at 30°C. Tryptone soya broth (TSB) and *Pseudomonas Isolation Agar* (PIA) were used for specific experiments. Antibiotics were used at (µg ml^-1^): rifampicin (Rif), 20 for *P. putida*; kanamycin (Km), 50 for *P. putida* and 25 for *E. coli*; ampicillin (Amp), 100 for *E. coli*; gentamycin (Gm), 25 for *P. putida* and 10 for *E. coli*; chloramphenicol (Cm), 10 for *E. coli;* nalidixic acid (Nal), 10 for *E. coli;* streptomycin (Sm), 400 for *P. putida* and 50 for *E. coli*.

### Construction of plasmids and bacterial strains

The plasmids and primers used in this study are listed in Tables S2 and S3, respectively. DNA manipulations were performed using standard methods (Sambrook *et al*., 1989). Q5® High-Fidelity DNA polymerase (New England Biolabs) was used for PCR reactions according to the manufacturer’s instructions. Primers were synthesised by Macrogen, Inc. and restriction enzymes were purchased from New England Biolabs. All DNA constructs were sequenced and verified before use. Recombinant plasmids were transferred to *E. coli* strains by transformation and to *Pseudomonas* strains by electroporation (Choi *et al*., 2006) or conjugation (Ramos-Gonzalez *et al*., 1991), as appropriate.

Translational fusions to *lacZ* were designed for genes containing putative Rsm binding sites in the 5’ untranslated regions or the first codons of the coding regions, namely *tagB1*, *tssE1*, *tssJ1*, *hcp1*, *hcp4* and *hcp5*. These fusions cover between 120-540 bps of the coding region (*i.e.* 40-180 codons) and ∼ 500 bps upstream of the starting codon to include any possible intergenic promoter region or with the inclusion of the artificial promoter *P_tac_* instead. The DNA containing the above-described regions was amplified from genomic DNA extracted from *P. putida* KT2440 using primers P1-P12 (Table S3). Promoters were cloned into the broad host range and low copy number vector pSEVA225T (Silva-Rocha et al., 2013; Martínez-García et al., 2023) at the XbaI or BamHI/HindIII sites to produce translational fusions to the *lacZ* gene.

*P. putida rsmIEA tssA1* was constructed by allelic exchange, as previously described (Vasseur et al., 2005), using *P. putida rsmIEA* (Huertas-Rosales et al., 2016) as the recipient strain. *P. putida gacS* and *rsmIEA gacS* were constructed using *P. putida* and *P. putida rsmIEA* as the recipient strains, respectively. Briefly, 500-bp DNA fragments upstream and downstream of the gene to be deleted (*tssA1 or gacS*) were amplified using *P. putida* KT2440 genomic DNA and primers P13-P16 (Table S3). A fragment containing both regions was obtained by overlapping PCR, cloned into pJET1.2/blunt (Thermo Scientific™), sequenced and subcloned into the XbaI/BamHI sites of pKNG101 to generate pKNG101-*tssA1* (Bernal et al., 2017) and pKNG101-*gacS*. The suicide vector pKNG101 (Kaniga et al., 1991) does not replicate in *Pseudomonas*; it was maintained in *E. coli* CC118λpir and mobilised into *Pseudomonas* by triparental conjugation. The deletions were confirmed by PCR and DNA sequencing (Macrogen Inc.).

A similar approach was used to replace wildtype *P. putida tke2* and *tke7* genes with versions encoding the protein of interest C-terminally fused to a double V5 tag (Table S2). Primers P17-P20 (Table S3) were used to engineer the *tke7* substitution, while the *tke2-V5* construct is described in (Bernal *et al*., 2017). The same strategy was used to replace wildtype *P. putida tssB1* with the gene encoding the protein of interest, C-terminally fused to sfGFP, as described in (Bernal *et al*., 2021). All insertions and gene replacements were confirmed by PCR and DNA sequencing.

Gene *hcp5* was amplified from genomic DNA extracted from *P. putida* KT2440 hcp5-StrepII using primers P21-P22 (Table S3) designed to fuse a StrepII tag sequence at the 3’end. The *hcp5* gene was cloned into the broad host range, medium copy number vector pSEVA234 (Table S2 (Martínez-García *et al*., 2023)) from the SEVA collection at the KpnI/XbaI sites to generate pSEVA234-hcp5::StrepII. Primers P23-P24 were used for screening and sequencing purposes.

Gene *hcp1* was amplified from genomic DNA extracted from *P. putida* KT2440 using the primers P25-P26 (Table S3). The *hcp1* gene was cloned into the bacterial two-hybrid vector pKT25 (Table S2, (Karimova *et al*., 1998)) at the XbaI/BamHI sites to produce pKT25-hcp1 which expresses a T25 fragment C-terminal fused to Hcp1 (T25-Hcp1). Primers P27-P28, which bind to the upstream and downstream pKT25 insertion sites, were used to screen colonies harbouring the *hcp1* insert and for subsequent sequencing.

Gene *hcp5* was amplified using primers P29-P30 (Table S3). The *hcp5* gene was cloned into the bacterial two-hybrid vector pUT18 (Table S2, (Karimova et al., 1998)) at the XbaI/BamHI sites to produce pUT18-hcp5 which expresses a T18 fragment N-terminal fused to Hcp5 (Hcp5-T18). Primers P31-P32 were used to screen colonies harbouring the *hcp5* insert and for subsequent sequencing.

The genes *tke7* and *tke2* were amplified from genomic DNA extracted from *P. putida* KT2440 using primers P33-P34 and PP37-38, respectively (Table S3). For tke7, primers P35-P36 encoding the *pelB* signal sequence were used to introduce *pelB* at the 5’ end of *tke7* at the NheI restriction site. This sequence encodes a 22 N-terminal leader peptide MKYLLPTAAAGLLLLAAQPAMA (UniProt Q04085) originally from the pectate lyase B (PelB) of *Erwinia carotovora* CE (Lei *et al*., 1987). PelB directs the fused protein, Tke7, to the periplasm via the Sec system pathway, where a signal peptidase removes the signal peptide (Tsirigotaki *et al*., 2017). The *tke7* and *tke2* genes were cloned into the broad host range, medium copy number vector pS238D•M (Table S2 (Calles et al., 2019)) from the SEVA collection at the NheI/BamHI sites, which excised the *msf*-GPF to produce pS238D•*pelB*-*tke7* and pS238D•*tke2*. Primers P23-P24, which bind to the upstream and downstream pS238D•M insertion sites, were used to screen colonies harbouring the *pelB-tke7* or *tke2* insert and for subsequent sequencing.

### Identification of Rsm binding sites

The *P. putida* KT2400 genome was obtained from the *Pseudomonas* Genome Database (NC_002947) (Winsor et al., 2016). To search for Rsm targets along the T6SS cluster sequences and the orphan genes (*hcp4-6* and *vgrG4-5*), we used the tool ‘find motifs’ in Geneious 2023.2.1, based on the EMBOSS v6.5.7 tools fuzznuc and fuzzpro, and the three motifs 5’-nnncanggangn-3’, 5’-nnnnnnggangn-3’ and 5’-nnncanggann-3’ that contain the GGA core and the most conserved bases of the consensus sequences 5′-RU<u>ACA</u>R<u>GGA</u>U<u>GU</u>-3′ (Dubey *et al*., 2005; Duss *et al*., 2014), and 5’-<u>^A^</u>/_U_<u>CA</u>N<u>GGA</u>NG^U^/_A_-3’ (Schubert *et al*., 2007). We searched the forward and reverse strands, allowing for zero mismatches. The 1308 hits were exported, and a Perl script (extract_annotations_with_CDS_direction.pl) was used to link the annotations from the KT2440 GenBank file with those from the hits table: perl extract_annotations_with_CDS_direction.pl > annotations_with_CDS_direction.txt. Finally, the table was converted to a tab-delimited format using the Python script: python convert_table.py. We manually curated these hits, eliminating those in the opposite direction of encoding genes. The outcome of this analysis is presented in Dataset S1, which is organised into one tab for each T6SS cluster or orphan gene.

The secondary RNA structure of the Rsm binding sites was predicted using Mfold (Zuker, 2003) with the default settings (Folding temperature is fixed to 37°C; ionic conditions: 1M NaCl, no divalent ions; linear RNA folding at 5%, window = 0, max folds = 50) and selecting the output structure with the lowest energy. The input sequence was 20-30 nucleotides long, including the Rsm-binding site sequence of 12 nucleotides and ∼4-9 nucleotides upstream and downstream of this region.

### ***β***-galactosidase assay

Overnight cultures of the *P. putida* strains carrying the pSEVA225T plasmid and their derivatives were diluted to a final OD_600_ of 0.05 in fresh LB medium containing kanamycin; and the cultures were grown at 30°C and 180 RPM for 4 h. OD_600_ reached aproximately 0.7 after 4 h and 5.5 after 24 h, and at this point, aliquots were taken to measure β-galactosidase activity in permeabilised whole cells as described by (Miller, 1972). At least five independent assays were performed for each case, and the standard errors of the mean were calculated for each.

### Secretion assay

*P. putida* strains were grown in TSB for 8 h at 30°C with shaking at 180 RPM, and the extracellular fraction was obtained and analysed as previously described (Bernal *et al*., 2021). Briefly, the cell suspensions were spun three times at 10,000 x g for 20 min, and the top fraction was collected each time. Bacterial pellets were normalised and added directly to 1x Laemmli buffer, while the culture supernatants were collected and precipitated with trichloroacetic acid (TCA) overnight. The precipitants were then washed with acetone, resuspended in 1x Laemmli buffer, and boiled for 15 min. SDS-PAGE analysis was performed using 4-20% Mini-PROTEAN® TGX Stain-Free^TM^ Precast Protein Gels (Bio-Rad), TGS/SDS running buffer (144 g/L of glycine, 30 g/L Tris and 1% SDS) and prestained protein ladder (bioBLU Prestained Protein Ladder, gTPbio). Proteins were transferred to a 0.2 µm PVDF membrane (Bio-Rad) using a Transfer Pack and a Trans-Blot Turbo transfer system (Bio-Rad) before blocking in 5% w/v skim milk/TBS-T and adding the primary and secondary antibodies. The following primary antibodies were used: rabbit anti-Hcp1 antibody (dilution 1:500 in 5 w/v % skim milk/TBS-T), mouse *E. coli* anti-RNA polymerase beta (BioLegend) (dilution 1:5,000 in 5 w/v % skim milk/TBS-T), mouse anti-V5 (Cell Signaling Technology) (dilution 1:1,000 in 5 w/v % skim milk/TBS-T) or Strep-Tactin-HRP/AP conjugate (Iba Lifesciences) (dilution 1:3,000 in 3 w/v % BSA/TBS-T) at 4°C overnight or room temperature for 4 h. The following secondary antibodies were used: goat anti-rabbit IgG-HRP conjugate (Cell Signaling Technology) (dilution 1:5,000 in 5% w/v skimmed milk/TBS-T) and horse anti-mouse IgG-HRP conjugate (Cell Signaling Technology) (dilution 1:5,000 in 5% w/v skimmed milk/TBS-T). The membranes were washed three times for 5 min with TBS-T prior to development. HRP conjugates were visualised with Immobilon Forte Western HRP substrate (Merck) using an Odyssey® Fc imager (LI-COR BioSciences) and the Li-COR® acquisition software. At least three independent experiments were conducted.

### Flow cytometry

Flow cytometry was used to monitor the expression of the translational GFP fusions. Data acquisition was performed using a Cytomics FC500-MPL cytometer (Beckman Coulter) and analysed with FlowJo X version 10.0.7r software (Tree Star, Inc.). Overnight LB-grown cultures of *P. putida* KT2440 expressing TssB1-sfGFP and the isogenic *rsmIEA* and *gacS* mutants were washed and diluted in PBS to a final concentration of 10^-7^ cells ml^-1^ for fluorescence measurement. A 2D histogram plotting side scatter (SSC) versus forward scatter (FSC) was used to distinguish bacterial cells from the background noise, with an FSC threshold value of 3. The bacterial cell population was gated in this histogram for subsequent measurements of GFP fluorescence intensity. Fluorescence values from 100,000 events were compared with those of a reporterless control strain to determine the fractions of ON and OFF cells. All experiments were performed in at least three independent replicates.

### Interbacterial competition assays

*In vitro* competition assays were performed on LB Lennox (5 g/L NaCl) agar plates (1.5% w/v), as previously described (Civantos *et al*., 2024). Briefly, overnight bacterial cultures were washed and adjusted to an OD_600_ of 10 in sterile phosphate-buffered saline (PBS) and mixed at a 1:1 ratio (*P. putida*:prey). The mixtures were grown on LB agar plates at 30°C for 5 h (*E. coli, Salmonella* and *Erwinia* prey), collected using an inoculating loop, and resuspended in sterile PBS. The outcome of the competition was quantified by counting colony-forming units (CFUs) using antibiotic selection of the input (time = 0 h) and output (time = 5 h). All prey strains harboured the plasmid pRL662, which confers resistance to gentamicin, whereas *P. putida* KT2440 harboured the plasmid pSEVA234, which confers resistance to kanamycin. For competition assays, at least five biologically independent experiments were conducted. The competitive index values were calculated using the following formula: competitive index = (output attacker/output prey)/(input attacker/input prey).

### Clustal Omega

The six Hcp proteins identified in the genome of *P. putida* KT2440 (Hcp1: PP3089, Hcp2: PP4082, Hcp3: PP2615, Hcp4: PP0655, Hcp5: PP4886 and Hcp6: PP5238) were aligned using the Multiple Sequence Alignment (MSA) tool of Clustal Omega (EMBL-EBI) (Madeira *et al*., 2024) with default settings.

### Statistical analyses

Statistical analyses are based on t-tests in which two conditions are compared independently. P-values from raw data were calculated by two-tailed t-test and from ratio data to the control by one-sample t-test using GraphPad Prism 8 version 8.3.0 and are represented in the graphs by ns, non-significant; *P < 0.05; **P < 0.01; and ***P < 0.001.

## RESULTS

### *P. putida* KT2440 T6SS transcripts present Rsm binding sites

To study the RsmIEA-dependent regulation of the different components of *P. putida* T6SS, we first identified putative binding sites along the sequence of the three clusters and the orphan genes *hcp4-6* and *vgrG4-5* (Bernal *et al*., 2017). These sites were identified based on similarity to the Rsm binding site consensus sequences 5′-RU<u>ACA</u>R<u>GGA</u>U<u>GU</u>-3′ (Dubey *et al*., 2005; Duss *et al*., 2014), and 5’-<u>^A^</u>/_U_<u>CA</u>N<u>GGA</u>NG^U^/_A_-3’ (Schubert *et al*., 2007) with a special focus on the presence of the GGA core. We identified a large number of putative binding sites for RsmIEA along the sequence of the three clusters and the orphan *hcp* and *vgrG* genes, with some regions exhibiting particularly high density (Dataset S1 and Fig. 1A). Next, we cross-referenced this information with the experimental data obtained in a genome-wide analysis of *P. putida* RNA targets post-transcriptionally regulated by Rsm proteins performed by <u>R</u>NA <u>A</u>ntisense <u>P</u>urification followed by <u>Seq</u>uencing (RAP-Seq, (Huertas-Rosales *et al*., 2021)). These data validated our previous predictions of Rsm binding sites, revealing that RsmI, RsmE and RsmA (hereafter referred to as RsmIEA) bind large regions of mRNA molecules encoding several T6SS elements of the K1-T6SS and the orphan clusters *hcp4* and *hcp5*. These regions included *tagB1-tssB1-tssC1-tssE1*, *clpV1-tssJ1-tssK1*, *hcp1-tssA1*, *tke2-tki2* and *hcp4* and *hcp5* mRNAs (Fig. 1A and Dataset S1).

**Fig. 1.**
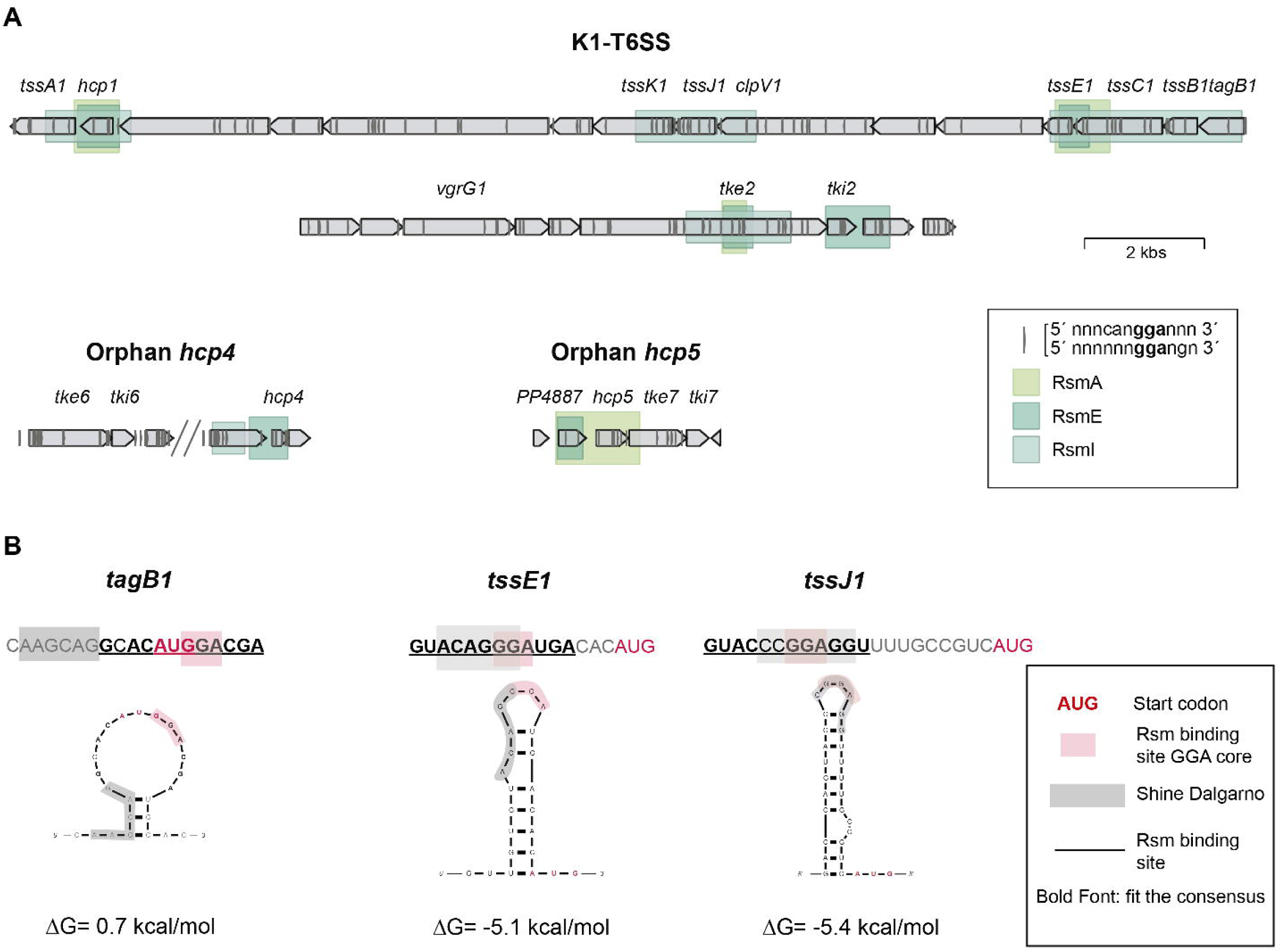
Rsm binding regions and sites identified in *P. putida* KT2440 T6SS transcripts. **A)** Green boxes depict Rsm protein binding regions in the K1-T6SS and orphan *hcp4* and *hcp5* as identified by (Huertas-Rosales *et al*., 2021). The vertical lines represent putative binding sites that fit the consensus sequence and are identified in the *in silico* study performed in this work, supporting likely direct binding to transcribed mRNAs from these regions. **B)** Secondary structure of the RNA targets (*tagB1*, *tssE1* and *tssJ1*) using Mfold software. The start codon is shown in red, the Rsm binding site GGA core is boxed in pink while the putative Shine Dalgarno is boxed in grey. The putative Rsm binding site sequences is underlined and the bases that fit the consensus sequence are in bold.

Consistently, Rsm binding sites were located near the RBS either blocking or promoting translation, fitting with their known binding preference. From the previously identified potential Rsm binding sites, we selectively focused on those near the start codon of the abovementioned T6SS transcripts, including *tagB1*, *tssE1* and *tssJ1* (Table S4), for further investigation. Interestingly, the sequences of three *hcp* mRNA molecules (*hcp1*, *hcp4* and *hcp5*) revealed multiple putative Rsm binding sites and were identified as RsmIEA targets in the RAP-Seq analysis (Fig. 1A, Table S4 and Dataset S1; Huertas-Rosales et al., 2021). These *hcp* RNA sequences were included in the subsequent assays.

First, to evaluate the quality of the selected Rsm binding sites, we predicted the secondary structure of the RNA targets using Mfold software. The RNA binding affinity is highly dependent on the secondary structure, with optimal interactions occurring when GGA is either in the loop of the hairpin or within it (Dubey *et al*., 2005). Our prediction indicated that the selected T6SS mRNA targets could form short stem-loop structures with the GGA core contained within the hairpin loop (Fig. 1B and S1). In all cases, *i.e. tagB1*, *tssE1* and *tssJ1 (hcp1, hcp4* and *hcp5)*, the GGA core overlapped with the Shine-Dalgarno sequence or the AUG start codon (Fig. 1B and S1). These data strongly support that Rsm proteins bind directly to these mRNA targets.

### RsmIEA repress the translation of T6SS transcripts

Since Rsm proteins, in most cases, interfere with ribosome binding and prevent translation of target transcripts, we expect to observe reduced translation of T6SS mRNA molecules targeted by Rsm proteins. To study the level of translation of these target genes, we generated ectopic translational fusions to the promoterless *lacZ* gene. This reporter gene encodes the β-galactosidase enzyme that converts a colourless substrate into a colourful substance, allowing it to measure the levels of translation colourimetrically (Miller, 1972). The constructs consist of either the native operon promoter or an introduced *P_tac_* promoter fused to *lacZ* alongside the first portion of the encoding region of each target gene.

Since the GacS/GacA two-component system regulates Rsm proteins, the translational fusions were introduced in wildtype, *gacS* and *rsmIEA* mutant strains. The level of translation of the selected T6SS transcripts showed a marked decrease in *gacS* and an increase in *rsmIEA* mutants (Fig. 2). LacZ activity of all these transcripts was severely decreased in the *gacS* mutant, with levels similar to the empty plasmid control (Fig. 2; blue and grey columns, respectively). Robust increases were observed for *tagB1, tssE1, hcp4* and *hcp5* and a 40-fold increase for *hcp1* and *tssJ1* when comparing the wildtype to the *rsmIEA* background (Fig. 2; green and red columns, respectively). This data demonstrates that RsmIEA proteins repress the translation of the K1-T6SS gene cluster and the orphan *hcp4* and *hcp5* mRNAs and this repression is blocked by the sRNAs that are positively governed by the GacS/GacA cascade.

**Figure 2.**
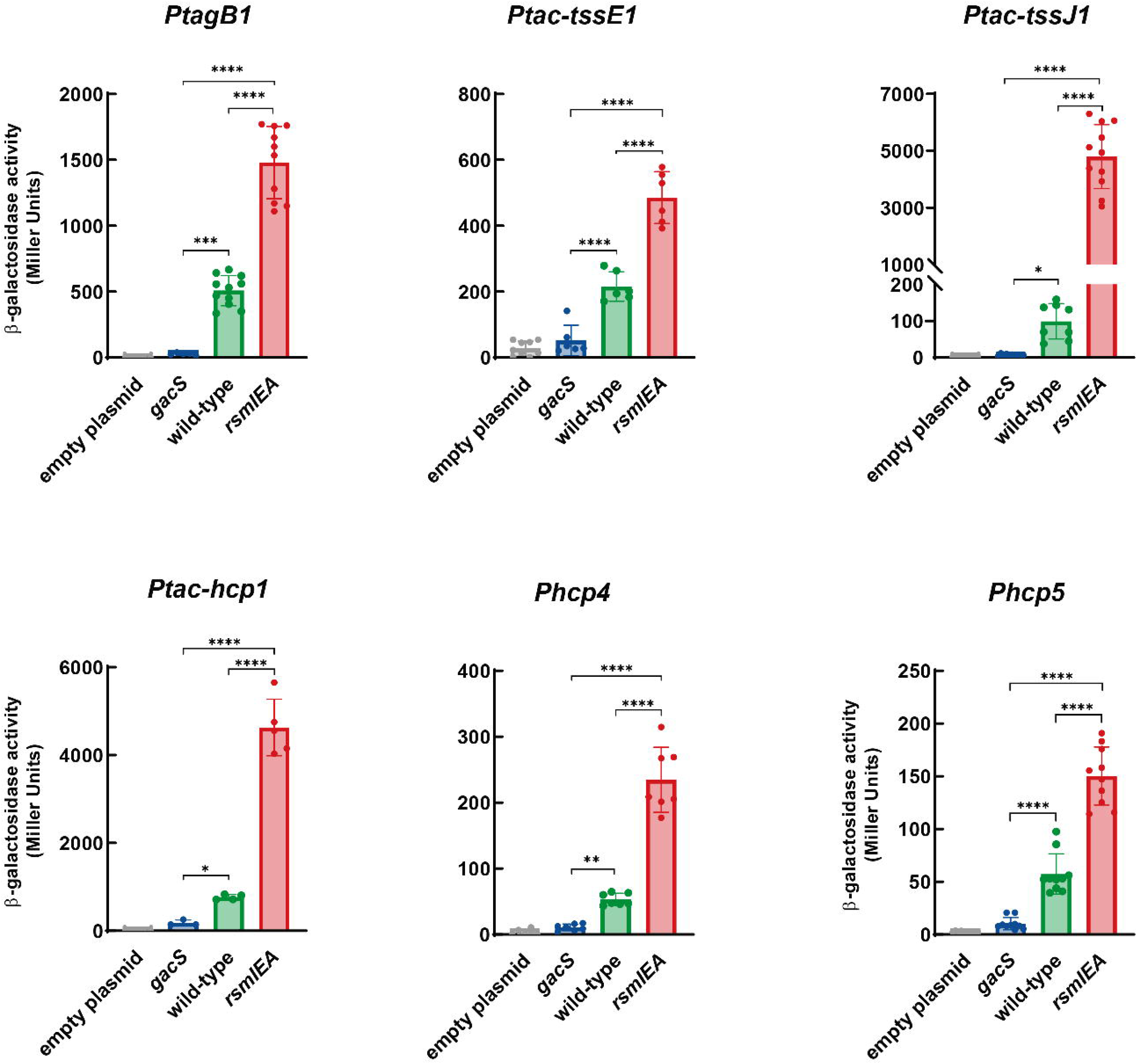
Deletion of *gacS* represses, whilst deletion of *rsmIEA* enhances translation of T6SS transcripts. **A**) β-galactosidase activity, in exponential phase, of the *P. putida* wildtype and the *gacS* or *rsmIEA* mutant strains bearing the pSEVA225T or the pSEVA225T-derivated plasmids containing the indicated translational fusions (P*_tagB1_*, P*_tssE1_*, P*_tssJ1_*, P*_hcp1_*, P*_hcp4_*, and P*_hcp5_*) to the *lacZ* gene. Strains were grown in LB, and the β-galactosidase activity was measured in the exponential phase of growth. Data are means ± SD from at least four replicates (*n* ≥4), each one including two technical replicates. P values were calculated by t-test analysis.

### RsmIEA represses the production and secretion of components of the K1-T6SS and the hcp5 orphan cluster

Next, we aimed to analyse whether the GacS and RsmIEA-dependent changes in the translational level of the T6SS transcripts (Fig. 2) are also converted into an increase in T6SS protein production and secretion.

Hcp proteins are a reliable marker to assess the functionality of T6SS since they assemble the internal tube that, upon contraction of the sheath, is ejected out of the cell when the system is active (Basler and Mekalanos, 2012). In *P. putida*, the detection of Hcp1 in the supernatant of wildtype cultures has been previously demonstrated (Bernal *et al*., 2017, 2021). This secretion is abolished in an isogenic *tssA1* mutant unable to encode the essential T6SS component TssA1 (Bernal *et al*., 2017, 2021). To compare the levels of production and secretion of Hcp1 between the wildtype, the *gacS* and the *rsmIEA* mutant strains, we used a polyclonal antibody to detect Hcp1. Using conditions we know activate the K1-T6SS, we observed highly repressed expression of Hcp1 in the *gacS* mutant but expression in both the WT and Rsm mutants (Fig. 3A, left panel). We found that the Hcp1 protein is overrepresented in the secretome in the *rsmIEA* mutant (Fig. 3A, left panel).

**Fig. 3.**
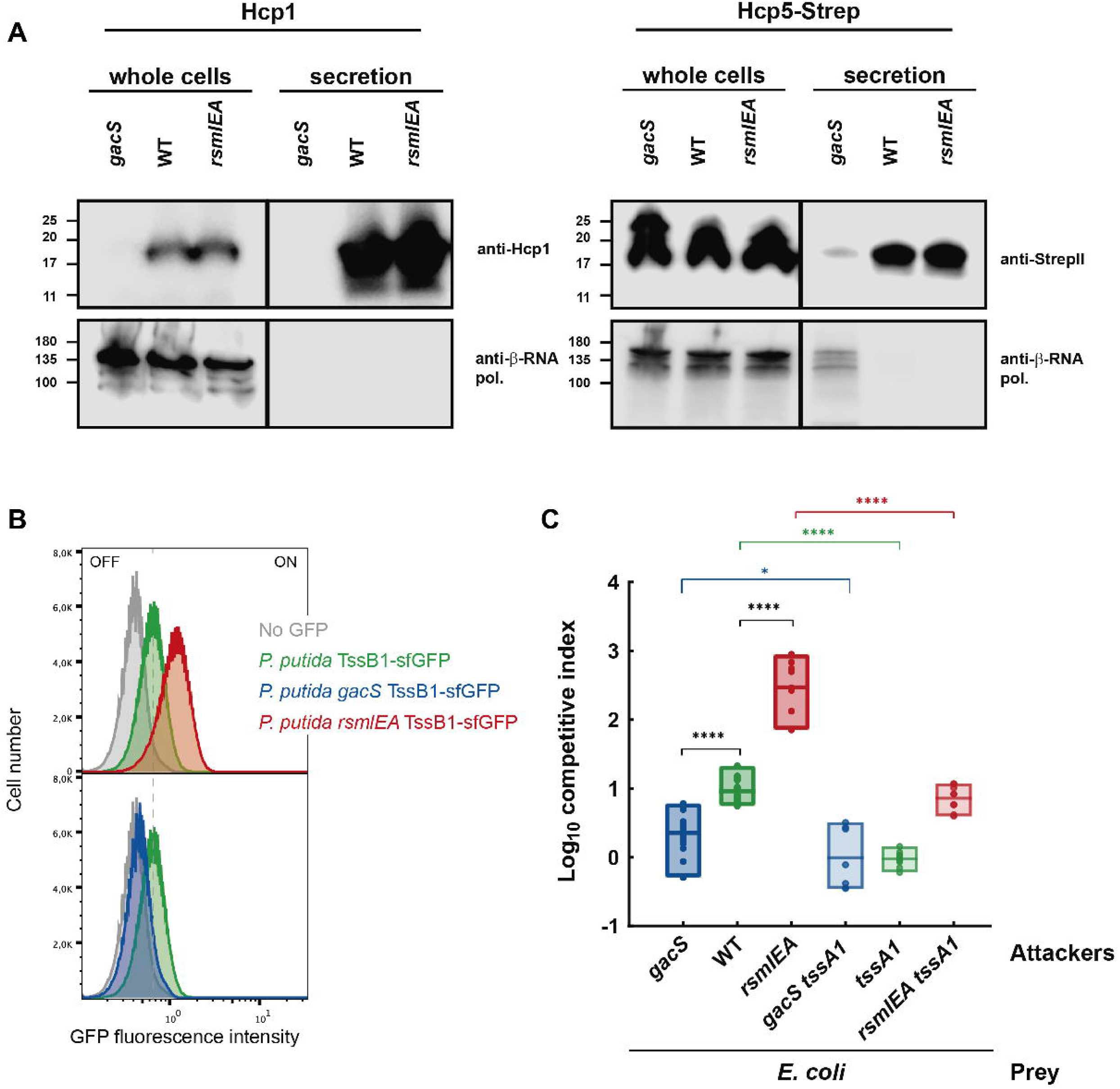
RsmIEA represses the production and secretion of components of the K1-T6SS and the *hcp5* orphan cluster. **A)** The *gacS* and the *rsmIEA* mutants of *P. putida* show markedly reduced and robust secretion of Hcp1 and Hcp5, respectively. Hcp1 and Hcp5-StrepII expression and presence in the culture supernatant were assessed for wildtype *P. putida* and the isogenic *gacS* and *rsmIEA* mutants. Hcp5-StrepII was expressed from plasmid pSEVA234 (Table S2). Positions of molecular weight markers are shown on the left, and the β-subunit of the *E. coli* RNA polymerase (β-RNA pol.) was used as a loading and bacterial lysis control. A representative blot from three independent experiments is presented. **B**) Representative flow cytometry profiles of *P. putida* wildtype and *gacS* and *rsmIEA* mutant strains demonstrate repressed or enhanced expression of TssB1-sfGFP from the native *tssB1* locus. **C**) Competition assays between *P. putida* strains and *E. coli* show a decreased competitive index for the *gacS* mutant compared to the wildtype *P. putida* strain. On the contrary, an increased competitive index is shown for the *rsmIEA* mutant compared to the wildtype *P. putida* strain. Graph shows means ±SD, significance is indicated by * = p < 0.05, ** = p < 0.01, *** = p < 0.001 or **** = p < 0.0001. Statistical analysis was performed using unpaired T-tests corrected for multiple testing using the Holm-Sidak method. For *E. coli* competitions, *n* = 5 (all significant).

To monitor the expression and secretion of an orphan Hcp, we selected Hcp5, as in KT2440, *hcp4* is a truncated gene (Bernal *et al*., 2017), and engineered a plasmid harbouring *hcp5* fused to the sequence encoding a twin StrepII tag. Although the expression of Hcp5 in the whole cell lysate in this condition is independent of the regulatory cascade (Fig. 3A, right panel), the secretion of Hcp5 likely requires an active K1-T6SS. These results confirmed the regulation of the K1-T6SS and their secreted components, such as the orphan Hcp5 protein, by GacS/GacA and Rsm systems.

While the western blot analysis provided a bulk-level quantification of total Hcp1 and Hcp5 protein, it averages out individual cellular behaviours and cannot distinguish whether the observed differences occur uniformly across the population. To resolve this phenotypic heterogeneity and precisely quantify the proportion of cells actively assembling the T6SS at the single-cell level, we performed flow cytometry, leveraging a TssB1-superfolder GFP (sfGFP) fusion expressed at its native locus. By tracking single-cell fluorescence to determine the proportion of TssB1-positive cells, we detected a heterogeneous population in the wildtype strain, with only ∼50–60% of the cells presenting TssB1-GFP, indicating T6SS assembly under the tested conditions (green, Fig. 3B). Notably, the *rsmIEA* mutant (red, Fig. 3B, Top) showed an increased population of cells containing an active K1-T6SS (∼80%) compared to the wildtype strain (green, Fig. 3B). In contrast, the *gacS* mutant showed no K1-T6SS activity (blue, Fig. 3B, Bottom), with a profile resembling the sfGFP-negative control (grey, Fig. 3B). Crucially, these single-cell insights reveal that the higher protein levels previously observed via western blot do not reflect an increased or faster secretion rate per individual cell. Instead, they demonstrate that these mutations alter population dynamics by recruiting a larger number of individual cells to actively assemble the system. These findings confirm that the post-transcriptional regulation of T6SS transcripts by the Gac and Rsm proteins dictates the precise percentage of active cells within the bacterial population.

Lastly, to determine if the increased number of K1-T6SS active cells would result in a higher capacity to kill, we investigated T6SS-dependent killing in this context. Consistent with the previous results, the *rsmIEA* mutant showed a significantly increased competitive index when cocultured with prey *E. coli* (CI^WT^ = 1 versus CI*^rsmIEA^* = 2.5; Fig. 3C). The increased killing ability of the *rsmIEA* mutant was, at least partially, K1-T6SS dependent, as the competitive index decreased in the isogenic *rsmIEA tssA1* mutant (CI*^rsmIEA^ ^tssA1^* = 0.86; Fig. 3C). In contrast, the *gacS* mutant presented a competitive index lower than both, the wildtype and the *rsmIEA* mutant (CI*^gacS^* = 0.4; Fig. 3C). This aligns with the inability of the *gacS* mutant to synthesise the small RNAs to sequester RsmIEA that would unleash the K1-T6SS activity.

Thus, the absence of RsmIEA proteins promotes the translation of K1-T6SS transcripts (Fig. 2), leading to the overproduction and secretion of specific K1-T6SS and orphan Hcp5 (Fig. 3A). This increases the percentage of the population assembling an active T6SS apparatus (Fig. 3B), ultimately enhancing the capacity to kill prey cells (Fig. 3C).

### Hcp5 is an additional component of the K1-T6SS

KT2440 possesses six Hcp proteins (Hcp1-6), three of them encoded within the K1-, K2- and K3-T6SS clusters and three orphan Hcp proteins (Hcp4, Hcp5 and Hcp6) encoded elsewhere in the genome. These orphan Hcp proteins are closely related to Hcp1, with an approximately 50% identity between Hcp1 and the orphan Hcp4, Hcp5 and Hcp6 (Fig. 4A). This homology suggests that the K1-T6SS secretes the orphan Hcp proteins. In contrast, Hcp2 and Hcp3, with a protein identity between them of 99.4% and ∼20% identity with Hcp1 and the orphans, clustered in a different phylogenetic group and are expected to be K2- and K3-T6SS-dependent, respectively (Fig. 4 A and B).

**Figure 4.**
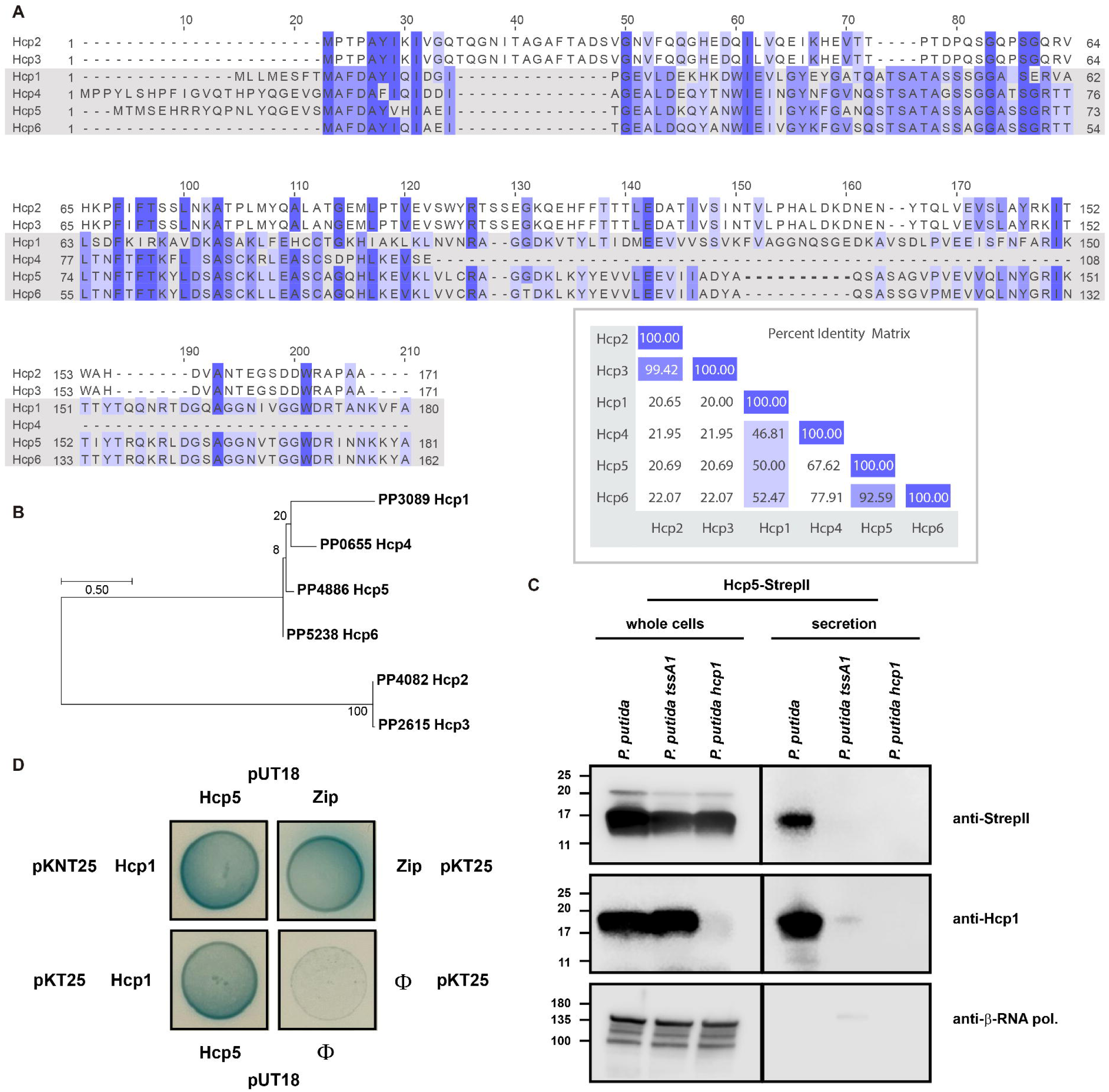
Functionality of the orphan Hcp5 protein. **A**) Multiple Sequence Alignment (MSA) of *P. putida* Hcp protein sequences using Clustal Omega and displayed by Jalview coloured by percentage of identity. The percent identity matrix generated by Clustal 2.1 is included in the bottom part of the figure. **B**) Phylogenetic tree of *P. putida* Hcp proteins. Maximum likelihood tree was built with Mega 7 software. **C**) Hcp5 expression and presence in the culture supernatant were assessed for wildtype *P. putida* and the isogenic *tssA1* and *hcp1* mutants. The *hcp1* mutant was used as a control for tube formation. A representative blot from three independent experiments is presented. **D**) *P. putida* Hcp5 interacts with Hcp1 by the N- and the C-terminal domains, as assessed by bacterial two-hybrid assay. Three independent experiments were performed, with identical results.

In other systems, such as *P. aeruginosa* T6SSs, orphan VgrG islands are co-regulated with the cluster to which they are associated (Allsopp *et al*., 2017). Here we show that *hcp4* and *hcp5* are co-regulated with the K1-T6SS by RsmIEA proteins (Figs. 1, 2 and 3). Thus, the homology and the co-regulation data support the idea that Hcp4 and Hcp5, are secreted through the K1-T6SS. To test our hypothesis, we introduced pSEVA234-*hcp5::StrepII* in both the wildtype and the *tssA1* mutant backgrounds. In the wildtype strain, we readily detected Hcp5 in the culture supernatant, but its secretion was abolished in the isogenic *tssA1* mutant (Fig. 4C), confirming that the secretion of Hcp5, like Hcp1, depends on the K1-T6SS.

We observed that in the *hcp1* mutant, Hcp5-StrepII is expressed, but the secretion of the orphan Hcp5 protein was abolished (Fig. 4C). To explore this idea further, we assessed if Hcp1 and Hcp5 could interact via a bacterial two-hybrid assay. We demonstrate pronounced blue X-gal metabolism, similar to our Zip-Zip positive control, demonstrating that Hcp1 and Hcp5 (Fig. 4D). These two pieces of data demonstrate that Hcp1 is the critical component for the assembly of the K1-T6SS inner tube and is required for a functional K1-T6SS. It also shows that Hcp1 and Hcp5 can directly interact, which suggests that the Hcp1-K1 tube can incorporate accessory Hcp proteins like Hcp5 that cannot be secreted in the absence of the main tube component, Hcp1 (Fig. 4C).

The incorporation of different Hcp proteins could serve as accessory proteins that facilitate the secretion of additional Hcp-associated effectors, thereby increasing the diversity of T6SS “bullets” deployed by the system.

### Tke2 and Tke7 are regulated by Rsm and GacA/S cascade

As early studies demonstrated, T6SS toxins are co-regulated alongside the clusters with which they are associated (Allsopp *et al*., 2017, 2022). In *P. putida*, toxins related to the RsmIEA-regulated K1-T6SS and *hcp5* orphan clusters, Tke2 and Tke7, respectively, seem to follow this pattern. The transcripts of *tke2* and *tke7* are partially bound to RsmIEA proteins (Fig. 1A), and putative Rsm binding sites are identified through their sequences (Fig. 1A). To compare the levels of production/secretion of these effectors between the wildtype, the *rsmIEA* and *gacS* mutant strains, we introduced a V5 tag at the C-terminus of the native copy of Tke2 and Tke7 in these backgrounds. Western blot analysis detected expression of both Tke2 and Tke7, which is slightly more abundant in the exponential phase of growth in the *rsmIEA* mutant and highly repressed in the *gacS* mutant compared to the wildtype strain (Fig. 5).

**Fig. 5.**
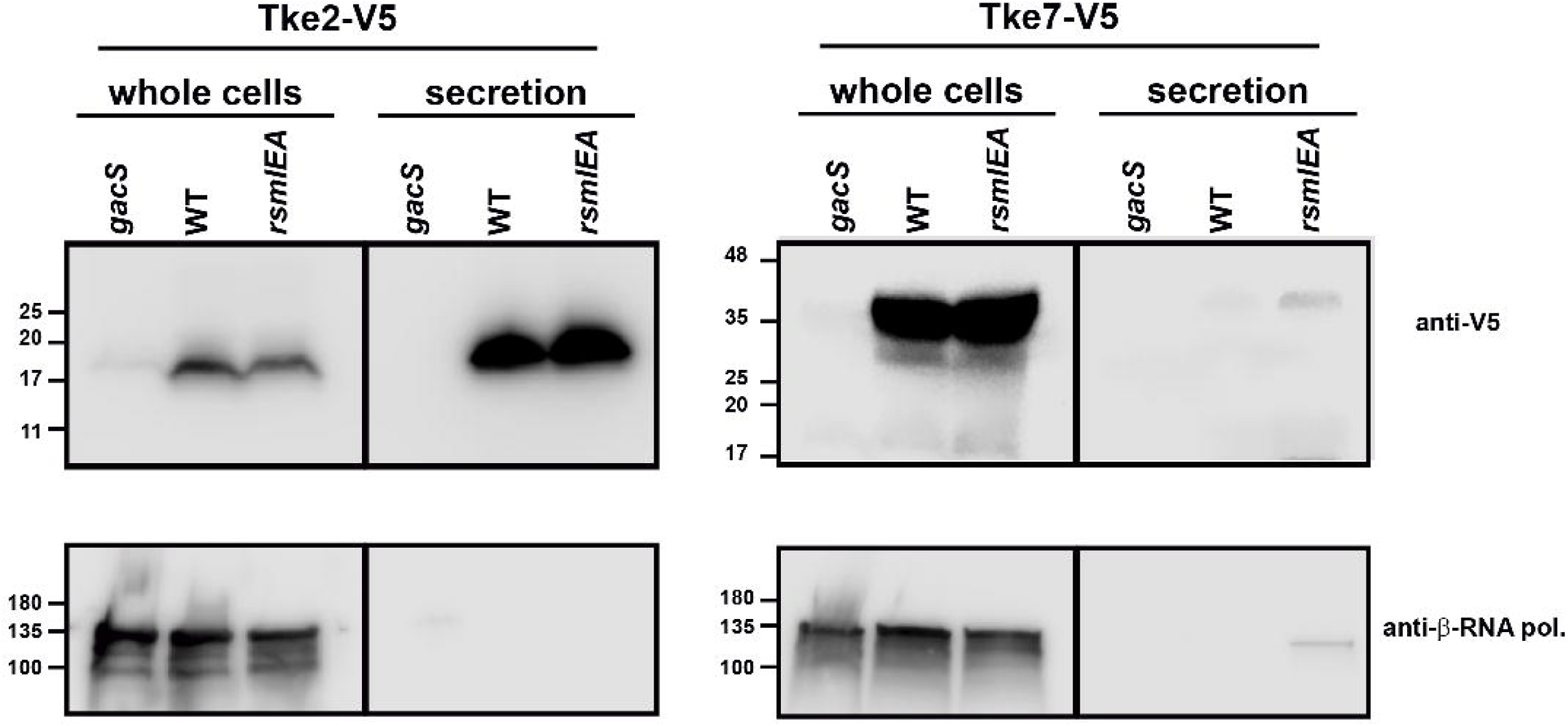
Tke2 and Tke7 are regulated by Rsm and GacS/A cascade, but only secretion of Tke2 was observed. Production and secretion of Tke2 and Tke7 effectors were assessed for wild-type *P. putida* and the isogenic *tssA1* mutant. A representative blot from three independent experiments is presented.

Given that the K1-T6SS is predominantly induced during the stationary phase of growth, we next sought to evaluate the processing and secretion dynamics of these effectors in this phase. Tke2 is an Rhs effector that can be detected in whole cells as both the full-length protein and the C-terminal domain (∼16 kDa) (Fig. 5 and Fig. S2). However, while both forms are present intracellularly, the C-terminal domain is the predominant form found in the secreted fraction. Crucially, this secretion is entirely absent in a structural *tssA1* mutant (Fig. S2), confirming T6SS-dependent secretion. Tke7 is likewise detected in whole cells (Fig. 5 and S2), but it was not detected in the secreted fraction under the tested conditions.

In summary, the absence of RsmIEA proteins promotes the translation of K1-T6SS-dependent effector transcripts (Fig. 2), leading to enhanced production of Tke2 and Tke7 effectors and secretion of specific K1-T6SS and *hcp5* orphan cluster components, including Hcp5 and Tke2 (Fig. 3A and 5).

### RsmIEA regulates the biocontrol capability of this strain by repressing Tke2 and Tke7 effectors

Since *P. putida* is a biocontrol agent and the T6SS is instrumental in the killing capacity of this strain, we decided to test whether the two RsmIEA-regulated effectors, Tke2 and Tke7 (Fig. 1A), are part of the *P. putida* arsenal to kill plant pathogens or other pathogens of importance in agriculture. To investigate the toxicity of Tke2 and Tke7, we selected a recalcitrant plant pathogen, *Erwinia amylovora,* the causal agent of a devastating necrotic disease (fire blight) affecting apples, pears, and other rosaceous plants and a human pathogen, *Salmonella enterica,* that produces salmonellosis (diarrhoea, nausea, vomiting, abdominal pain) if consumed (Malnoy *et al*., 2012; Galán, 2021). We heterologously expressed the effectors from pS238•DM and performed growth curves. Since Tke7 is predicted to be a pore-forming toxin, we added a PelB sequence to send the effector to the periplasm as described before for Tke5 (Velázquez, Arce-Rodríguez, *et al*., 2026). Upon induction with 3-*m*Bz, the viability of the pathogens was severely decreased when both toxins were expressed in comparison with the non-induced controls and the vector expressing *msfGFP* (Fig. 6A, 6B and S3). Notably, Tke2 exhibits a stronger effect on *Salmonella* while Tke7 is more effective against *Erwinia*, suggesting a certain degree of target specificity (Fig. 6A, 6B and S3). Collectively, these experiments demonstrate the potent antibacterial activity of the Tke2 and Tke7 toxins against environmentally found pathogens. This inhibitory potential indicates that Tke2 and Tke7 contribute to the biocontrol capabilities of *Pseudomonas putida* in agricultural settings, offering promising tools to target pathogens that threaten the food industry.

**Fig. 6.**
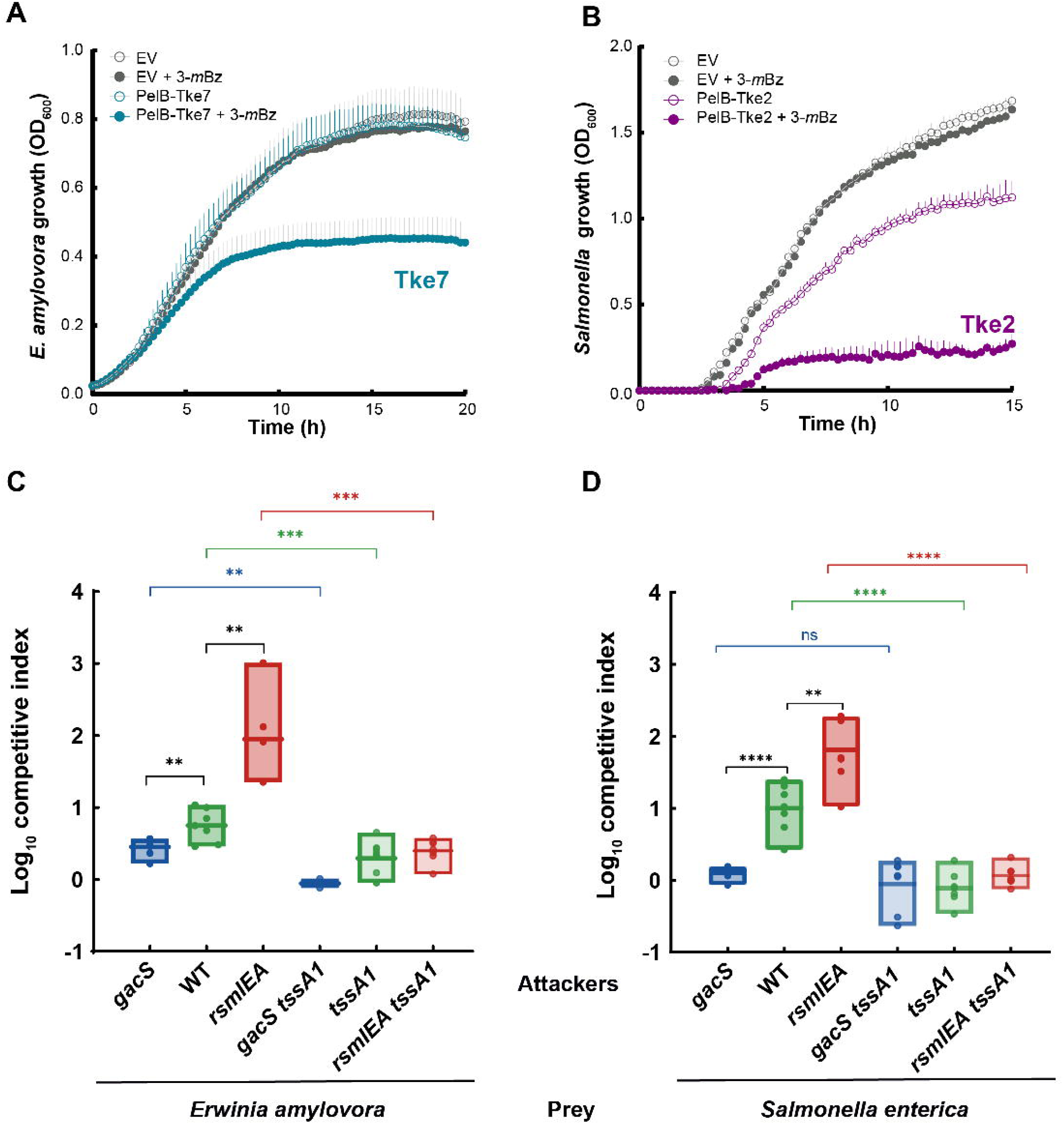
Toxicity of Tke2 and Tke7 for different recalcitrant pathogens. The growth of **A**) the plant pathogen *Erwinia amilovora* cells harbouring the pS238D•*pelB*-*tke7* containing the T6SS effector of *P. putida* Tke7 N-terminal fused to a PelB signal peptide and **B**) the human pathogen *Salmonella enterica* subsp. enterica serovar Senftenberg cells harbouring the pS238D•*tke2* containing the T6SS effector of *P. putida* Tke2 were determined by measuring the OD at 600 nm. After 2 h, 3-*m*Bz was added to the LB medium to induce the expression of the *tke2* and *pelB-tke7* gene. **C**) and **D**) Competition assays between *P. putida* strains, *Salmonella*, and *Erwinia* show an increased competitive index for the *rsmIEA* mutant and a decreased competitive index for the *gacS* mutant compared to the wildtype *P. putida* strain. A positive log_10_ C.I. indicates the attacker successfully outcompeted the prey, while a value of zero signifies equal bacterial fitness. Data are mean ± SD. *P < 0.05; **P < 0.01; ***P < 0.001; ****P < 0.0001. Statistical analysis was performed using unpaired t tests corrected for multiple testing using the Holm–Sidak method. For *Salmonella* comparisons, *n* = 5, for *Erwinia* comparisons, *n* = 5.

Since the effectors regulated by these proteins present antimicrobial activity against *E. amylovora* and *Salmonella*, we decided to test the ability of the *gacS* and *rsmIEA* mutants to eliminate these pathogens. Importantly, we observed that the *rsmIEA* mutant showed an enhanced capacity to kill *E. amylovora* (CI^WT^ = 0.75 versus CI*^rsmIEA^* = 2; Fig. 6C) and *Salmonella* (CI^WT^= 0.9 versus CI*^rsmIEA^* = 1.8; Fig. 6D). In contrast, the *gacS* mutant, unable to sequester RsmIEA proteins so has a repressed K1-T6SS, presented a lower competitive index that the wildtype strain in both cases, against *E. amylovora* (CI^WT^ = 0.75 versus CI*^gacS^* = 0.45; Fig. 6C) and *Salmonella* (CI^WT^ = 1 versus CI*^gacS^* = 0.1; Fig. 6C). These data demonstrated the capacity of *P. putida* to kill pathogens of interest for agriculture and human health via the T6SS.

We conclude that the K1-T6SS and its associated effector proteins represent a powerful biocontrol mechanism, with significant potential for agricultural applications. Understanding the regulatory networks governing its expression and characterising its secreted antimicrobial arsenal will enhance its effectiveness as a targeted biocontrol tool in crop protection strategies.

## DISCUSSION

The T6SS is widely recognised as a bacterial nanoweapon that delivers toxic effectors into neighbouring cells to gain a competitive advantage in polymicrobial environments (Allsopp and Bernal, 2023; Vázquez-Arias *et al*., 2025). The well-established biocontrol agent *P. putida* KT2440 deploys its T6SS to eradicate a broad spectrum of resilient phytopathogens, including *Xanthomonas campestris* and *Agrobacterium tumefaciens*, thereby protecting plants from infection (Bernal *et al*., 2017, 2021; Vázquez-Arias *et al*., 2025; Velázquez, Arce-Rodríguez, *et al*., 2026). This highlights the immense potential of *P. putida* T6SSs in biotechnological applications for sustainable agriculture.

The activity of T6SSs is controlled at multiple levels, including transcriptional, post-transcriptional, and post-translational mechanisms, ensuring their precise deployment when needed (Hespanhol *et al*., 2023). While the transcriptional regulation of the *P. putida* K1-T6SS has revealed constitutive basal expression that is further induced in the stationary phase and influenced by global regulators such as RpoN, RpoS, FleQ, RetS, and the GacS/GacA two-component system (Bernal *et al*., 2023), the post-transcriptional and post-translational control in this strain has remained poorly understood.

This study demonstrates that the RsmIEA proteins in *P. putida* serve as key post-transcriptional regulators of the K1-T6SS and associated orphan elements (Fig. 7). This regulatory function is achieved by RsmIEA binding to specific sites near the ribosomal binding sites (RBS) of T6SS transcripts, including those encoding Hcp1 and Hcp5, thereby repressing their translation (Fig. 1 and 7). This mechanism is analogous to the well-characterised RsmA/CsrA family of RNA-binding proteins in *Pseudomonas aeruginosa*, which also suppresses T6SS mRNA translation (Allsopp *et al*., 2017). The integration of RsmIEA-mediated post-transcriptional control with the established transcriptional regulatory networks (such as the GacS/GacA cascade) highlights the layered complexity that enables *P. putida* to fine-tune T6SS expression, assembly, and secretion. This intricate regulation is crucial for a likely energetically costly system, ensuring its optimal deployment in competitive environmental niches like the rhizosphere.

**Fig. 7.**
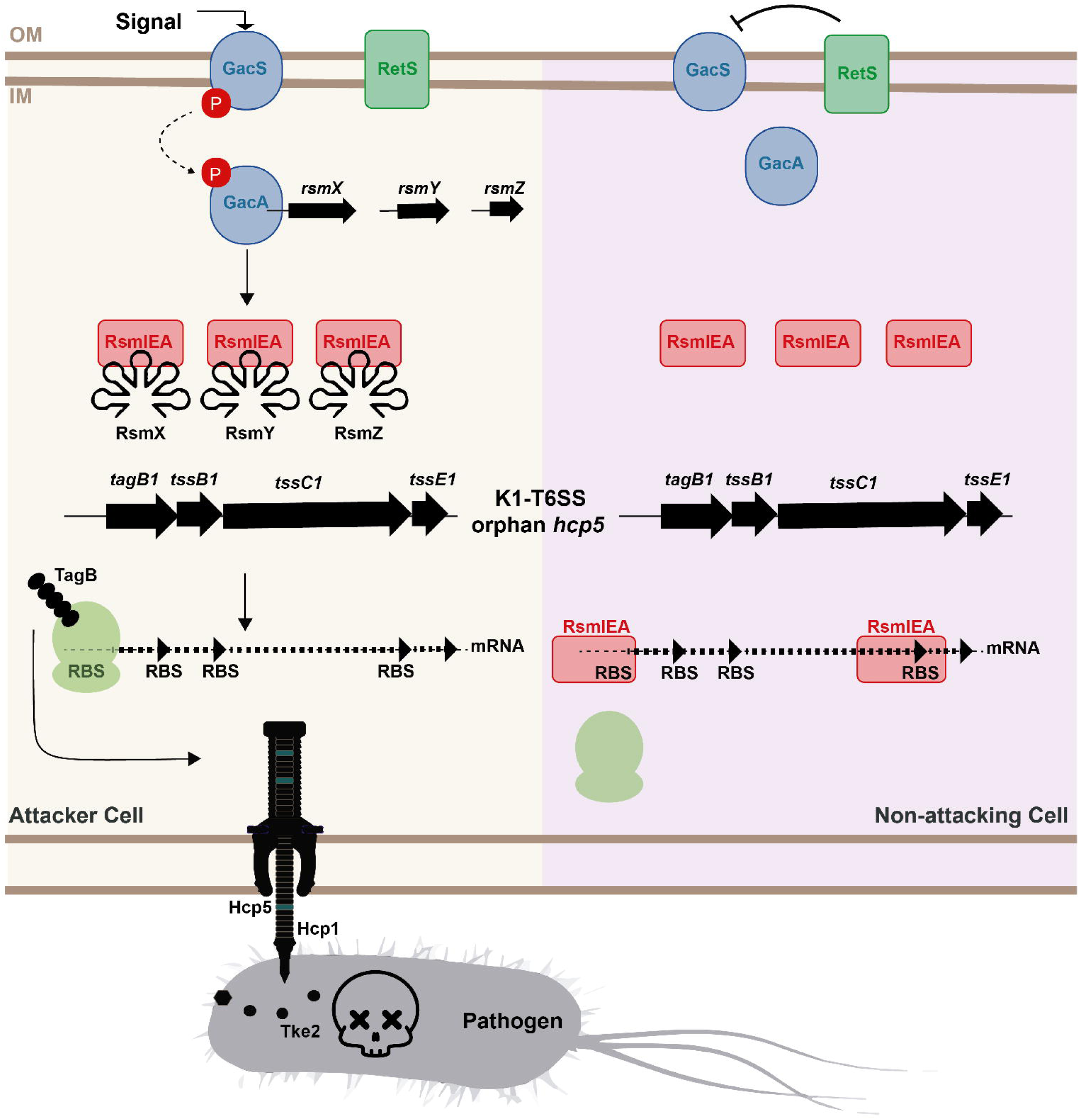
Model of the Gac/Rsm-mediated regulation of the K1-T6SS and the orphan *hcp5* in *P. putida*. The figure illustrates the regulatory switch determining the transition between an attacker and a non-attacking cell. In the attacker cell state, depicted on the left panel, an environmental signal triggers the autophosphorylation of the sensor kinase GacS, which transfers its phosphate group to the response regulator GacA. Activated GacA-P drives the transcription of the regulatory small RNAs RsmX, RsmY, and RsmZ, which subsequently sequester the translational repressors RsmIEA. This sequestration, or similarly, a *rsmIEA* mutant, leaves the ribosome-binding sites of the K1-T6SS and orphan *hcp5* mRNAs accessible, allowing ribosomes to bind, initiate translation, and produce the T6SS components like TagB. As a result, the active T6SS apparatus is assembled, enabling the cell to fire and kill competing pathogens using effectors like Tke2. Conversely, in the non-attacking cell state shown on the right panel, the absence of the signal or the inhibitory action of RetS on GacS, or a *gacS* mutant strain, leaves GacA unphosphorylated, halting the transcription of the small RNAs. The free RsmIEA proteins are then able to bind directly to the specific ribosome-binding sites on the polycistronic transcripts, causing steric hindrance that blocks ribosome access and prevents translation. Consequently, the T6SS nanoweapon cannot be assembled, allowing the bacterium to conserve metabolic energy.

The major limitation of our study is that the *rsmIEA* triple mutant was used, meaning the individual contributions of specific Rsm proteins were not assessed. This was a deliberate strategic choice, given that autoregulation and cross-regulation are well-documented within these systems. Additionally, ectopic expression from a plasmid encoding a single *rsm* gene could result in non-physiological expression levels. Nevertheless, our use of both *gacS* and *rsm* mutants clearly demonstrates the involvement of this pathway in regulating these T6SS genes. Indeed, Figure 1 shows that specific Rsm proteins exhibit binding preferences for distinct mRNAs, suggesting more nuanced levels of control.

A notable finding is the co-regulation of orphan T6SS components, particularly the *hcp5* cluster and its associated effector Tke7, by the GacS/A and RsmIEA systems (Fig. 2 and 3). *P. putida* KT2440 has a diverse T6SS arsenal, including three T6SS clusters (K1-, K2-, K3-T6SS) and five orphan gene clusters encoding Hcp and VgrG proteins with their respective effectors (Bernal *et al*., 2017). These orphan genes, likely acquired via horizontal gene transfer, significantly enhance the versatility of the T6SS, enabling the delivery of a broad array of effectors. Here, we identified Hcp1 as the essential component for the K1-T6SS inner tube assembly (Fig. 4 and 7). Our study indicates that the orphan Hcp5 can be incorporated into the Hcp1-K1 tube, potentially facilitating the secretion of additional effectors, such as Tke7. Recently, mixed tubes formed by hetero-hexamers with variable stoichiometry of minor and major Hcp complexes have been described in Bacteroidales (San-Miguel *et al*., 2026). The authors hypothesise that minor subunits act as recognition particles for effectors like Bte1 to mediate secretion. This is analogous to what we believe is happening, where Hcp5 likely directs Tke7 to the Hcp1-Hcp5 lumen tube. This modular structure allows the T6SS to deliver a diverse payload of effector proteins, broadening its efficacy across different target organisms.

The identification of Tke7 as a novel K1-T6SS effector (Fig. 6) further expands the known weaponry of *P. putida*. Tke7, which is predicted to be a pore-forming effector dependent on Hcp5 for its secretion, adds to the growing list of diverse effectors found in *P. putida*, which also includes the Rhs-type nuclease Tke2 and the pore-forming colicin Tke5 (Bernal *et al*., 2017; Velázquez, Arce-Rodríguez, *et al*., 2026; Velázquez, Zabala-Zearreta, *et al*., 2026). Tke5, for instance, has been shown to effectively kill a wide range of phytopathogens, including *P. syringae* and *Ralstonia solanacearum*, by disrupting bacterial membranes through selective ion transport (Velázquez, Arce-Rodríguez, *et al*., 2026). The discovery of such novel toxins with distinct mechanisms of action is critical, as they present promising therapeutic options against multidrug-resistant bacteria and phytopathogens. The existence of trans-kingdom effectors, capable of targeting bacteria, fungi, and eukaryotic cells, further broadens the potential application of these toxins.

In conclusion, this research provides crucial insights into the post-transcriptional regulation of the *P. putida* T6SS by RsmIEA proteins (Fig. 7) and enriches our understanding of its effector repertoire with the identification of Tke7. Optimising T6SS activity through a detailed comprehension of its multi-layered regulation directly enhances the potential of *P. putida* as an effective biocontrol agent. By leveraging this knowledge, it may be possible to engineer “designer microbes” with precisely controlled T6SSs, leading to more targeted and potent strategies for combating plant diseases and shaping the rhizosphere microbiota for greener agriculture. Future research should continue to explore the intricate regulatory cues, the full diversity of T6SS effectors and the mechanisms of action to fully harness these remarkable bacterial nanomachines for sustainable agricultural solutions and beyond.

## ACKNOWLEDGEMENTS

We thank María Milagros López and Joaquín Bernal Bayard for their kind gifts of *Erwinia* and *Salmonella* strains, respectively.

## DATA AVAILABILITY STATEMENT

All data generated during this study that support the findings are included in the manuscript or the Supplementary data. Uncropped and unedited gel images are included in Supplementary Fig. 4. The scripts for linking EMBOSS fuzznuc/fuzzpro motif-search hits to CDS annotations in *Pseudomonas putida* KT2440 have been deposited in Zenodo (https://doi.org/10.5281/zenodo.21257859).

## AUTHOR CONTRIBUTIONS (CRediT)

Conceptualisation: PB

Data curation: PB

Formal analysis: PB, JB

Funding acquisition: PB

Investigation: CC, CP, MM-T, MO, JB

Methodology: CC, CP, MM-T, MO, JB

Project administration: PB

Resources: PB

Supervision: MS-R, LPA, PB

Visualisation: DV-A, MS-R, PB

Writing - original draft: PB

Writing - review & editing: MS-R, LPA, PB

## REFERENCES

1. Allsopp, L.P. and Bernal, P. (2023) Killing in the name of: T6SS structure and effector diversity. Microbiology (N Y) 169: 1367.

2. Allsopp, L.P., Wood, T.E., Howard, S.A., Maggiorelli, F., Nolan, L.M., Wettstadt, S., and Filloux, A. (2017) RsmA and AmrZ orchestrate the assembly of all three type VI secretion systems in *Pseudomonas aeruginosa*. Proc Natl Acad Sci U S A 114: 7707–7712.

3. Arnold, T., Zeth, K., and Linke, D. (2009) Structure and function of colicin S4, a colicin with a duplicated receptor-binding domain. Journal of Biological Chemistry 284: 6403–6413.

4. Basler, M. and Mekalanos, J.J. (2012) Type 6 secretion dynamics within and between bacterial cells. Science (1979) 337: 815.

5. Bernal, P., Allsopp, L.P., Filloux, A., and Llamas, M.A. (2017) The *Pseudomonas putida* T6SS is a plant warden against phytopathogens. ISME Journal 11: 972–987.

6. Bernal, P., Civantos, C., Pacheco-Sánchez, D., Quesada, J.M., Filloux, A., and Llamas, M.A. (2023) Transcriptional organization and regulation of the *Pseudomonas putida* K1 type VI secretion system gene cluster. Microbiology (N Y) 169: 001295.

7. Bernal, P., Furniss, R.C.D., Fecht, S., Leung, R.C.Y., Spiga, L., Mavridou, D.A.I., and Filloux, A. (2021) A novel stabilization mechanism for the type VI secretion system sheath. Proc Natl Acad Sci U S A 118: e2008500118.

8. Bernal, P., Llamas, M.A., and Filloux, A. (2018) Type VI secretion systems in plant-associated bacteria. Environ Microbiol 20: 1–15.

9. Bernard, C.S., Brunet, Y.R., Gavioli, M., Lloubès, R., and Cascales, E. (2011) Regulation of type VI Secretion gene clusters by σ54 and cognate enhancer binding proteins. J Bacteriol 193: 2158–2167.

10. Calles, B., Goñi-Moreno, Á., and Lorenzo, V. (2019) Digitalizing heterologous gene expression in Gram[negative bacteria with a portable ON/OFF module. Mol Syst Biol 15: e8777.

11. Choi, K.-H., Kumar, A., and Schweizer, H.P. (2006) A 10-min method for preparation of highly electrocompetent *Pseudomonas aeruginosa* cells: application for DNA fragment transfer between chromosomes and plasmid transformation. J Microbiol Methods 64: 391–397.

12. Civantos, C., Ruiz, A., and Bernal, P. (2024) A Robust Method to Perform *In Vitro* and *In Planta* Interbacterial Competition Assays: Killing Plant Pathogens by a Potent Biocontrol Agent. In Host-Pathogen Interactions: Methods and Protocols. Medina, C. and López-Baena, F.J. (eds). New York, NY: Springer US, pp. 115–129.

13. Dubey, A.K., Baker, C.S., Romeo, T., and Babitzke, P. (2005) RNA sequence and secondary structure participate in high-affinity CsrA-RNA interaction. RNA 11: 1579–1587.

14. Duss, O., Michel, E., Dit Konté, N.D., Schubert, M., and Allain, F.H.T. (2014) Molecular basis for the wide range of affinity found in Csr/Rsm protein–RNA recognition. Nucleic Acids Res 42: 5332–5346.

15. Hespanhol, J.T., Nóbrega-Silva, L., and Bayer-Santos, E. (2023) Regulation of type VI secretion systems at the transcriptional, posttranscriptional and posttranslational level. Microbiology (N Y) 169: 1376.

16. Huertas-Rosales, Ó., Ramos-González, M.I., and Espinosa-Urgel, M. (2016) Selfregulation and interplay of Rsm family proteins modulate the lifestyle of *Pseudomonas putida*. Appl Environ Microbiol 82: 5673–5686.

17. Huertas-Rosales, Ó., Romero, M., Chan, K.-G., Hong, K.-W., Cámara, M., Heeb, S., et al. (2021) Genome-Wide Analysis of Targets for Post-Transcriptional Regulation by Rsm Proteins in *Pseudomonas putida*. Front Mol Biosci 8: 624061.

18. Huertas-Rosales, O., Romero, M., Heeb, S., Espinosa-Urgel, M., Amara, M.C., and Ramos-Gonz Alez, I. (2017) The *Pseudomonas putida* CsrA/RsmA homologues negatively affect c-di-GMP pools and biofilm formation through the GGDEF/EAL response regulator CfcR. Environ Microbiol 19: 3551–3566.

19. Jumper, J., Evans, R., Pritzel, A., Green, T., Figurnov, M., Ronneberger, O., et al. (2021) Highly accurate protein structure prediction with AlphaFold. Nature 596: 583–589.

20. Kaniga, K., Delor, I., and Cornelis, G.R. (1991) A wide-host-range suicide vector for improving reverse genetics in Gram-negative bacteria: inactivation of the blaA gene of Yersinia enterocolitica. Gene 109: 137–141.

21. Kay, E., Dubuis, C., Haas, D., and Lindow, S.E. (2005) Three small RNAs jointly ensure secondary metabolism and biocontrol in Pseudomonas fluorescens CHA0. Proc Natl Acad Sci U S A 102: 17136–17141.

22. Kelley, L.A., Mezulis, S., Yates, C.M., Wass, M.N., and Sternberg, M.J.E. (2015) The Phyre2 web portal for protein modeling, prediction and analysis. Nat Protoc 10: 845–858.

23. Lei, S.-P., Lin, H.-C., Wang, S.-S., Callaway, J., and Wilcox, G. (1987) Characterization of the *Erwinia carotovora pelB* Gene and Its Product Pectate Lyase. J Bacteriol 169: 4379–4383.

24. Ma, J., Campbell, A., and Karlin, S. (2002) Correlations between Shine-Dalgarno sequences and gene features such as predicted expression levels and operon structures. J Bacteriol 184: 5733–5745.

25. Madeira, F., Madhusoodanan, N., Lee, J., Eusebi, A., Niewielska, A., Tivey, A.R.N., et al. (2024) The EMBL-EBI Job Dispatcher sequence analysis tools framework in 2024. Nucleic Acids Res 52: W521–W525.

26. Martínez-García, E., Fraile, S., Algar, E., Aparicio, T., Velázquez, E., Calles, B., et al. (2023) SEVA 4.0: an update of the Standard European Vector Architecture database for advanced analysis and programming of bacterial phenotypes. Nucleic Acids Res 51: D1558–D1567.

27. Meng, E.C., Goddard, T.D., Pettersen, E.F., Couch, G.S., Pearson, Z.J., Morris, J.H., and Ferrin, T.E. (2023) UCSF ChimeraX: Tools for structure building and analysis. Protein Science 32: e4792.

28. Miller, J.H. (1972) Experiments in molecular genetics., NY: Cold Spring Harbor Laboratory.

29. Mirdita, M., Schütze, K., Moriwaki, Y., Heo, L., Ovchinnikov, S., and Steinegger, M. (2022) ColabFold: making protein folding accessible to all. Nat Methods 19: 679–682.

30. Nikel, P.I., Benedetti, I., Wirth, N.T., de Lorenzo, V., and Calles, B. (2022) Standardization of regulatory nodes for engineering heterologous gene expression: a feasibility study. Microb Biotechnol 15: 2250–2265.

31. Ramos-Gonzalez, M.I., Duque, E., and Ramos, J.L. (1991) Conjugational transfer of recombinant DNA in cultures and in soils: host range of *Pseudomonas putida* TOL plasmids. Appl Environ Microbiol 57: 3020–3027.

32. Reimmann, C., Valverde, C., Kay, E., and Haas, D. (2005) Posttranscriptional repression of GacS/GacA-controlled genes by the RNA-binding protein RsmE acting together with RsmA in the biocontrol strain Pseudomonas fluorescens CHA0. J Bacteriol 187: 276–285.

33. Sambrook, J., Fritsch, E.F., and Maniatis, T. (1989) Molecular Cloning: A Laboratory Manual., 2nd edition. Cold Spring Harbor Laboratory (ed) New York: Cold Spring Harbor Laboratory Press.

34. Sana, T.G., Hachani, A., Bucior, I., Soscia, C., Garvis, S., Termine, E., et al. (2012) The second type VI secretion system of *Pseudomonas aeruginosa* strain PAO1 is regulated by quorum sensing and fur and modulates internalization in epithelial cells. Journal of Biological Chemistry 287: 27095–27105.

35. San-Miguel, S.G., Al-Ammari, M.K.S., Johansson, E., Kowalska, A., Sauer, U., Batista, P.R., et al. (2025) Bacteroidales T6SS minor Hcp subunits form heteromers recognising effectors. BioRxiv.

36. Schubert, M., Lapouge, K., Duss, O., Oberstrass, F.C., Jelesarov, I., Haas, D., and Allain, F.H.T. (2007) Molecular basis of messenger RNA recognition by the specific bacterial repressing clamp RsmA/CsrA. Nature Structural & Molecular Biology 2007 14:9 14: 807–813.

37. Shine, J. and Dalgarno, L. (1974) The 3’-Terminal Sequence of Escherichia coli 16S Ribosomal RNA: Complementarity to Nonsense Triplets and Ribosome Binding Sites (terminal labeling/stepwise degradation/protein synthesis/suppression).

38. Silva-Rocha, R., Martínez-García, E., Calles, B., Chavarría, M., Arce-Rodríguez, A., Heras, A. de las, et al. (2013) The Standard European Vector Architecture (SEVA): a coherent platform for the analysis and deployment of complex prokaryotic phenotypes. Nucleic Acids Res 41: D666–D675.

39. Storey, D., McNally, A., Åstrand, M., Santos, J.S.-P.G., Rodriguez-Escudero, I., Elmore, B., et al. (2020) *Klebsiella pneumoniae* type VI secretion system-mediated microbial competition is PhoPQ controlled and reactive oxygen species dependent. PLoS Pathog 16: e1007969.

40. Tsirigotaki, A., De Geyter, J., Šoštarić, N., Economou, A., and Karamanou, S. (2017) Protein export through the bacterial Sec pathway. Nat Rev Microbiol 15: 21–36.

41. Vakulskas, C.A., Potts, A.H., Babitzke, P., Ahmer, B.M.M., and Romeo, T. (2015) Regulation of Bacterial Virulence by Csr (Rsm) Systems. Microbiology and Molecular Biology Reviews 79: 193–224.

42. Vasseur, P., Vallet-Gely, I., Soscia, C., Genin, S., and Filloux, A. (2005) The pel genes of the *Pseudomonas aeruginosa* PAK strain are involved at early and late stages of biofilm formation. Microbiology (N Y) 151: 985–997.

43. Vázquez-Arias, D., Civantos, C., Durán-Wendt, D., Ruiz, A., Rivilla, R., Martín, M., and Bernal, P. (2025) The *Pseudomonas putida* Type VI Secretion Systems Shape the Tomato Rhizosphere Microbiota. ISME Communications.

44. Velázquez, C., Arce-Rodríguez, A., Altuna-Alvarez, J., Rojas-Palomino, J., Flores-Cerón, A., Ruiz, A., et al. (2025) Tke5 is a novel *Pseudomonas putida* toxin that depolarises membranes killing plant pathogens. BioARchive.

45. Weller, D.M. (2007) *Pseudomonas* Biocontrol Agents of Soilborne Pathogens: Looking Back Over 30 Years. Phytopathology 97: 250–256.

46. Winsor, G.L., Griffiths, E.J., Lo, R., Dhillon, B.K., Shay, J.A., and Brinkman, F.S.L. (2016) Enhanced annotations and features for comparing thousands of Pseudomonasgenomes in the Pseudomonas genome database. Nucleic Acids Res 44: D646–D653.

47. Wu, C.F., Lin, J.S., Shaw, G.C., and Lai, E.M. (2012) Acid-Induced Type VI Secretion System Is Regulated by ExoR-ChvG/ChvI Signaling Cascade in *Agrobacterium tumefaciens*. PLoS Pathog 8: e1002938.

48. Yusupova, G.Z., Yusupov, M.M., Cate, J.H.D., and Noller, H.F. (2001) The path of messenger RNA through the ribosome. Cell 106: 233–241.

49. Zuker, M. (2003) Mfold web server for nucleic acid folding and hybridization prediction. Nucleic Acids Res 31: 3406–3415.

