## Supplementary data for "Rsm-mediated post-translational control of the *Pseudomonas putida* Type VI Secretion System"

**This file includes:**

Figures S1 to S4

Tables S1 to S4

Legend for Dataset S1

Supplementary references

### SUPPLEMENTARY INFORMATION FIGURES

### Figure S1

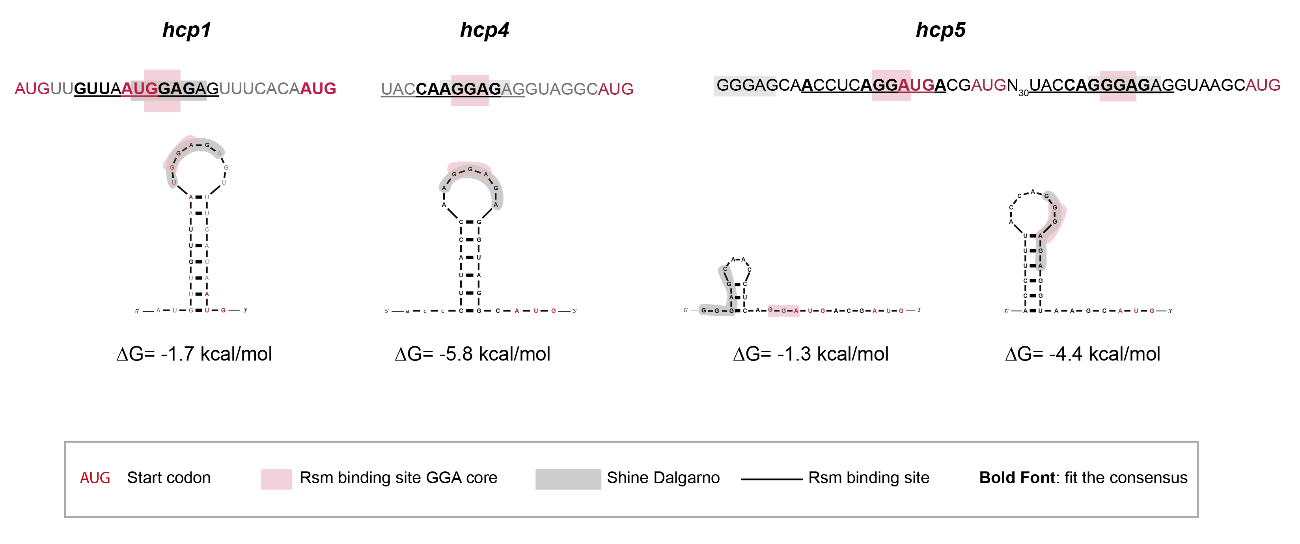

**Fig. S1. Rsm binding sites in *P. putida* KT2440 T6SS transcripts. A)** Secondary structure of the RNA targets (*hcp1*, *hcp4* and *hcp5*) using Mfold software. The start codon is shown in red, the Rsm binding site GGA core is boxed in pink, while the putative Shine-Dalgarno is boxed in grey. The Rsm binding site sequence, as identified in the *in silico* study, is underlined, and the bases that fit the consensus sequence are in bold.

### Figure S2

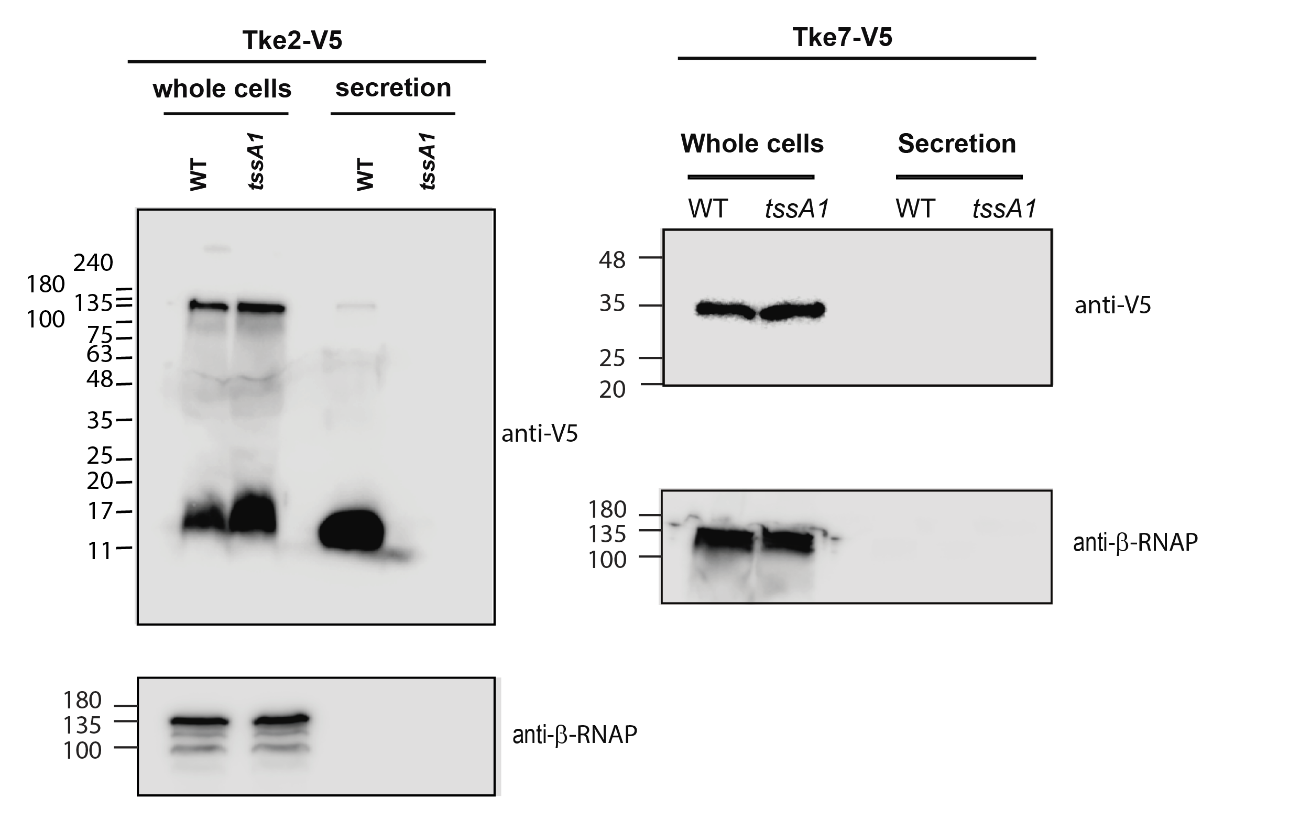

**Fig. S2. Production and secretion of Tke2 and Tke7 effectors. A)** The wildtype and the isogenic *tssA1* mutant of *P. putida* show production of Tke2-V5 in whole cells. As expected, Tke2-V5 is only present in the culture supernatant of the wildtype *P. putida* and absent in the *tssA1* mutant. Positions of molecular weight markers are shown on the left, the β-subunit of the *E. coli* RNA polymerase (β-RNA pol.) was used as a loading and bacterial lysis control and black lines indicate where the membrane was cut. A representative blot from three independent experiments is presented. **B**) Hcp5-StrepII was expressed from plasmid pSEVA234 (Table S2). Production and secretion of Tke7-V5 was assessed in the wildtype and the isogenic *tssA1* mutant of *P. putida.* While Tke7-V5 is present in whole cells, this effector is not detected in the supernatant of any of these strains. Positions of molecular weight markers are shown on the left, the β-subunit of the *E. coli* RNA polymerase (β-RNA pol.) was used as a loading and bacterial lysis control and black lines indicate where the membrane was cut. A representative blot from three independent experiments is presented.

### Figure S3

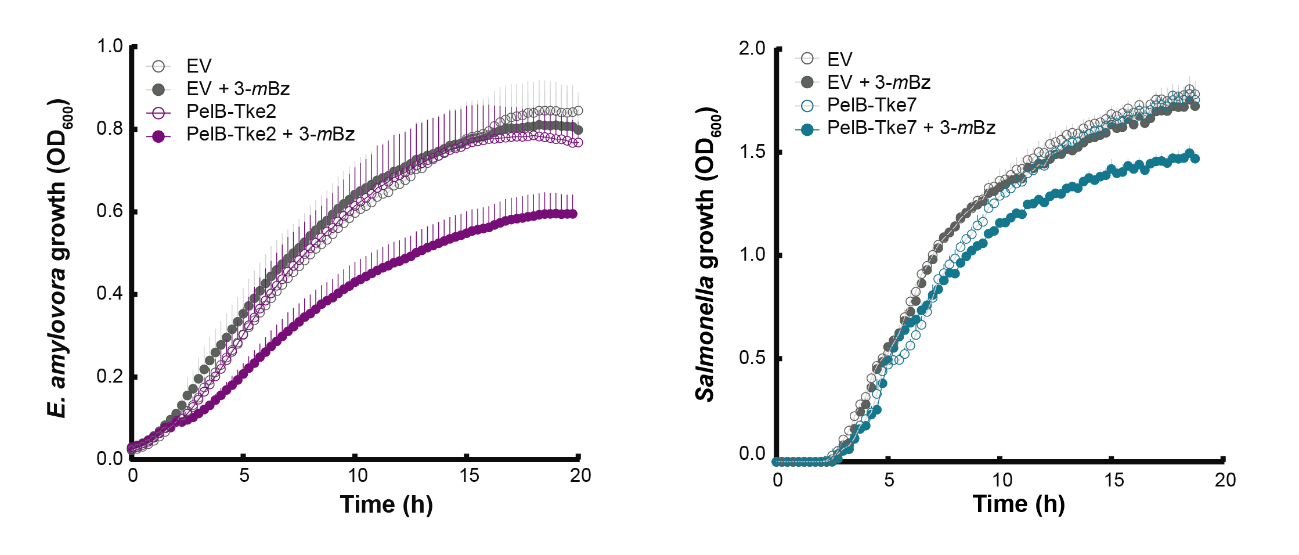

**Fig. S3. Toxicity of Tke2 and Tke7 for different recalcitrant pathogens.** The growth of **A**) the plant pathogen *Erwinia amilovora* cells harbouring the pS238D•*tke2* containing the T6SS effector of *P. putida* Tke2 and **B**) the human pathogen *Salmonella enterica* subsp. enterica serovar Senftenberg cells harbouring the pS238D•*pelB*-*tke7* containing the T6SS effector of *P. putida* Tke7 N-terminal fused to a PelB signal peptide were determined by measuring the OD at 600. After 2 hours, 3-*m*Bz was added to the LB medium to induce the expression of the *tke2* and *pelB-tke7* genes.

### Figure S4

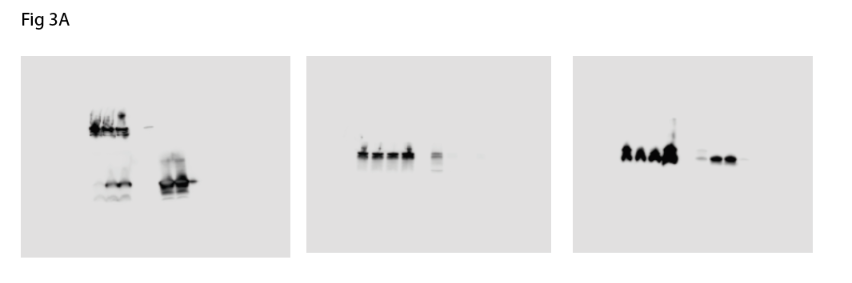

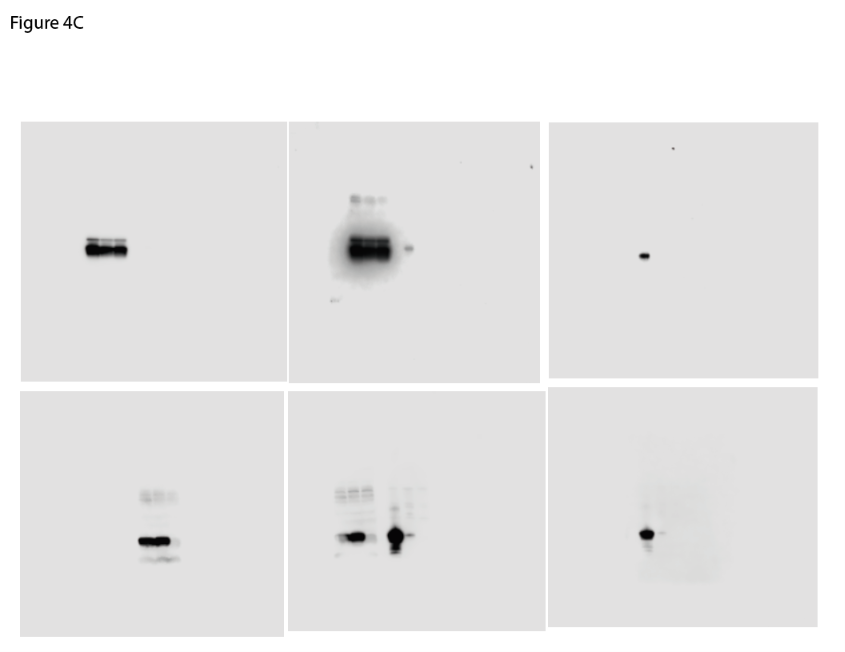

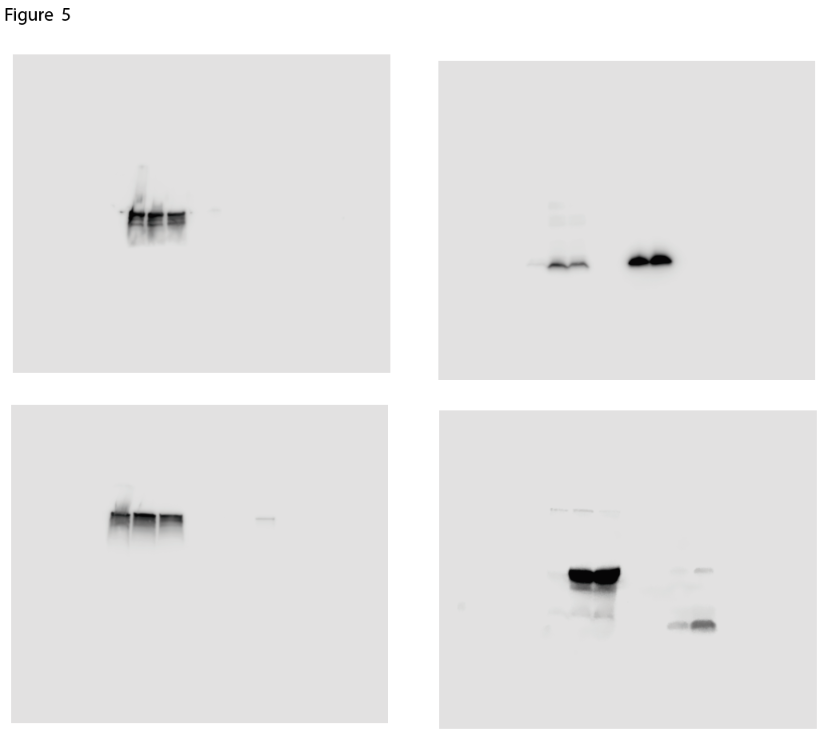

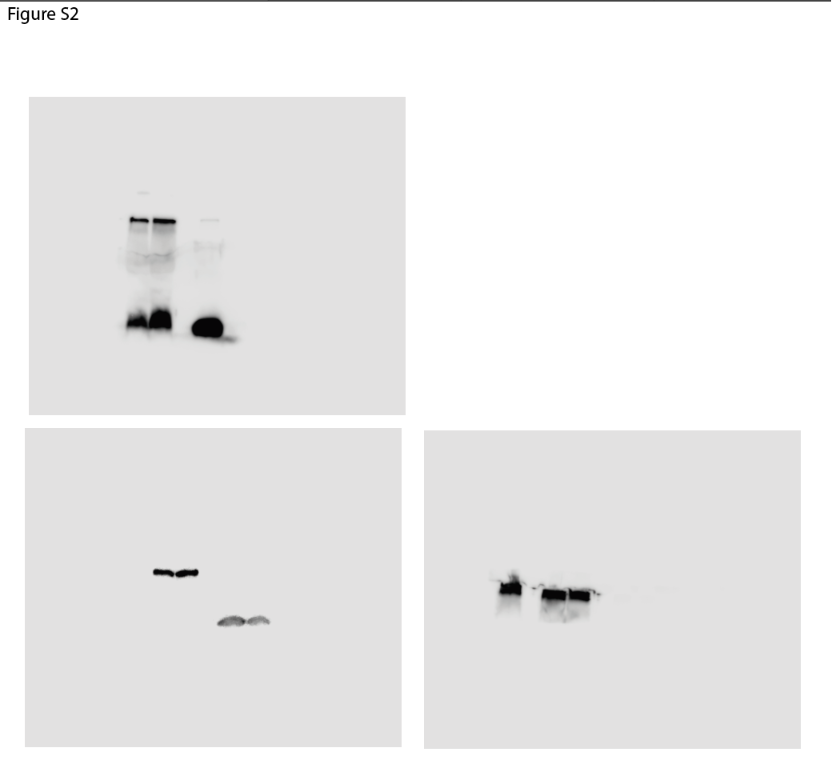

**Fig. S4.** Uncropped western blot membranes from Fig. 3, 4, 5 and Supplementary Fig. 2

### SUPPLEMENTARY INFORMATION TABLES

#### Table S1. Bacterial strains used in this study.

| **Name** | **Description** | | **Source** |
| --- | --- | --- | --- |
| ***Escherichia coli*** | | | |
| DH5α | F^–^ *end*A1 *gln*V44 *thi*-1 *rec*A1 *rel*A1 *gyr*A96 *deo*R *nup*G *pur*B20 φ80d*lacZ*∆M15 ∆(*lac*ZYA*-arg*F*)*U169 *hsd*R17(r_K_^–^m_K_^+^) λ^–^; Nal^R^ | | (Hanahan, 1985) |
| CC118λpir | *ara*D Δ(*ara*, *leu*) Δ*lac*Z74 *pho*A20 *gal*K *thi*-1 *rsp*E *rpo*B *arg*E *rec*A1 λ*pir* | | (Herrero *et al.*, 1990) |
| HB101 | supE44 hsdS20 recA13 ara-14 proA2 lacY1 galK2 rpsL20 xyl-5 mtl-1 | | (Boyer and Roulland-Dussoix, 1969) |
| ***Pseudomonas putida*** | | | |
| KT2440 | Wild-type, prototroph, cured of pWWO derivative of *P. putida* mt-2 | | (Nakazawa, 2002) |
| KT2440 *tssA1* | Markerless mutant | | This study |
| KT2440R *hcp1* | Markerless mutant, Rif^R^ | | (Bernal *et al.*, 2021) |
| KT2440 *rsmIEA* | Markerless mutant | | (Huertas-Rosales *et al.*, 2017) |
| KT2440 *gacS* | Markerless mutant | | This study |
| KT2440 *rsmIEA tssA1* | Markerless mutant | | This study |
| KT2440 *gacS tssA1* | Markerless mutant | | This study |
| KT2440 *tke2::V5* | Markerless mutant | | This study |
| KT2440 *rsmIEA* *tke2::V5* | Markerless mutant | | This study |
| KT2440 *gacS* *tke2::V5* | Markerless mutant | | This study |
| KT2440 *tssA1* *tke2::V5* | Markerless mutant | | This study |
| KT2440 *tke7::V5* | Markerless mutant | | This study |
| KT2440 *rsmIEA* *tke7::V5* | Markerless mutant | | This study |
| KT2440 *gacS tke7::V5* | Markerless mutant | | This study |
| KT2440 *tssA1* *tke2::V5* | Markerless mutant | | This study |
| KT2440 *tssB1::sfgfp* | Markerless mutant | | This study |
| KT2440 *rsmIEA::sfgfp* | Markerless mutant | | This study |
| KT2440 *gacS tssB1*::*sfgfp* | Markerless mutant, Rif^R^ | | This study |
| ***Plant and human pathogens*** | | | |
| *Erwinia amylovora* NCPPB 595 (CECT 222) | | CECT (Spanish Collection of Type Cultures) | |
| *Salmonella enterica sv. Senftenberg* | | Casadesus´ lab collection  Department of Genetics, University of Seville | |

#### Table S2. Plasmids used in this study. The antibiotic resistance markers are identified as follows: Amp, ampicillin; Km, kanamycin; Gm, gentamicin; Sm, streptomycin; Cm, chloramphenicol.

| **Name** | **Description** | **Source** |
| --- | --- | --- |
| pJET1.2/blunt | Cloning vector, ColE1 *ori*, Amp^R^ | Thermo Scientific |
| pSEVA225T | Standard vector used to construct *lacZ* translational fusion, RK2 *ori*, *'lacZ*, Km^R^ | (Martínez-García *et al.*, 2023) |
| pSEVA225T-P*tagB1* | *P. putida* P*_tagB1_* promoter region plus 120 bps of the 5´encoding sequence cloned into pSEVA225T as a translational fusion to *lacZ*, *P_tagB1_::lacZ*, Km^R^ | This study |
| pSEVA225T-Ptac-*tssE1* | *P_tac_* promoter fused to *P. putida tssE1* upstream region plus 120 bps of the 5´encoding sequence cloned into pSEVA225T as a translational fusion to *lacZ*, *P_tac-tssE1_::lacZ*, Km^R^ | This study |
| pSEVA225T-Ptac-*tssJ1* | *P_tac_* promoter fused to *P. putida tssJ1* upstream region plus 225 bps of the 5´encoding sequence cloned into pSEVA225T as a translational fusion to *lacZ*, *P_tac-tssJ1_::lacZ*, Km^R^ | This study |
| pSEVA225T-P*tac*-*hcp1* | *P_tac_* promoter fused to *P. putida tssJ1* upstream region plus 399 bps of the 5´encoding sequence cloned into pSEVA225T as a translational fusion to *lacZ*, *P_tac-hcp_1::lacZ*, Km^R^ | This study |
| pSEVA225T-*hcp4* | *P. putida* P*_hcp4_* promoter region plus 315 bps of the 5´encoding sequence cloned into pSEVA225T as a translational fusion, P*_hcp4_::lacZ*, Km^R^ | This study |
| pSEVA225T-*hcp5* | *P. putida* P*_hcp5_* promoter region plus 312 bps of the 5´encoding sequence cloned into pSEVA225T as a translational fusion, P*_hcp4_::lacZ*, Km^R^ | This study |
| pKNG101 | Gene replacement suicide vector, R6K *ori*, *sacB*, Sm^R^ | (Kaniga *et al.*, 1991) |
| pKNG101-*gacS* | PCR fragment containing _˜_550 bps region upstream and downstream of *gacS* gene cloned in pKNG101, to construct a clean deletion mutant of *gacS* gene containing only the 9 first and last base pairs of the *gacS* gene, Sm^R^ | This study |
| pKNG101-*tssB1::sfgfp* | PCR fragment containing the 3´ region of *tssB1* fused to *sfgfp* and the downstream region of the same gene cloned in pKNG101; when inserted into the chromosome and the plasmid cured, the strain expresses TssB1 C-terminally fused to sfGFP, Sm^R^ | Bernal et al 2021 |
| pKNG101-*tke2::V5* | PCR fragment containing the 3´ region of *tke2* fused to a 2x V5 tag and the downstream region of the same gene cloned in pKNG101, when inserted into the chromosome and the plasmid cured, the strain expresses Tke2^PP^ C-terminally fused to a 2xV5 tag, Str^R^ | Bernal et al 2017 |
| pKNG101-*tke7::V5* | PCR fragment containing the 3´ region of *tke7* fused to a 2x V5 tag and the downstream region of the same gene cloned in pKNG101, when inserted into the chromosome and the plasmid cured, the strain expresses Tke7 C-terminally fused to a 2xV5 tag, Str^R^ | This study |
| pRK600 | Helper plasmid, ColE1 ori, mobRK2, traRK2, Cam^R^ | (Kessler *et al.*, 1992) |
| pRL662-*gfp2* | pRL662 (Annette C. Vergunst *et al.*, 2000) derivative expressing GFP2, Gm^R^ | Erh-Min Lai lab |
| pSEVA234 | Expression vector; oriV(pBBR1) lacI^Q^ P_trc_, Km^R^ | (Martínez-García *et al.*, 2023) |
| pSEVA234-*hcp5::StrepII* | *hcp5* encoding Hcp5^PP^ with a C-terminal StrepII tag cloned into pSEVA234, pBBR1 *ori*, Km^R^ | This study |
| pS238D•M | Expression vector; oriV(pBBR1) XylS P_m_, lacI^Q^ P_trc_, Km^R^ | (Calles *et al.*, 2019) |
| pS238D•*pelB-tke7* | *tke7* encoding Tke7 with an N-terminal PelB leader sequence cloned into pS238DM*,* pBBR1 *ori*, Km^R^ | This study |
| pS238D•*tke2V5* | 3`end of *tke2* encoding the C-terminal domain (residues 1244-1385) of Tke2 cloned into pS238DM*,* pBBR1 *ori*, Km^R^ | This study |
| pKT25 | Bacterial-two-hybrid vector encoding the T25 fragment of *Bordetella pertussis* adenylate cyclase to construct C-terminal fusions, Km^R^ | (Karimova *et al.*, 1998) |
| pKT25-zip | T25 fragment of *Bordetella pertussis* adenylate cyclase C-terminally fused to Zip, Km^R^ | (Karimova *et al.*, 1998) |
| pKT25-hcp1 | T25 fragment of *Bordetella pertussis* adenylate cyclase C-terminally fused to Hcp1*^PP^*, Km^R^ | This study |
| pKNT25-hcp1 | T25 fragment of *Bordetella pertussis* adenylate cyclase N-terminally fused to Hcp1*^PP^*, Km^R^ | (Bernal *et al.*, 2021) |
| pUT18 | Bacterial-two-hybrid vector encoding the T18 fragment of *Bordetella pertussis* adenylate cyclase to construct N-terminal fusions, Amp^R^ | (Karimova *et al.*, 1998) |
| pUT18-zip | T18 fragment of *Bordetella pertussis* adenylate cyclase N-terminally fused to Zip, Amp^R^ | (Karimova *et al.*, 1998) |
| pUT18-hcp5 | T18 fragment of *Bordetella pertussis* adenylate cyclase N-terminally fused to Hcp5, Amp^R^ | This study |

#### Table S3. Oligonucleotide primers used in this study. The “Brief description” column provides basic information on the primer design (restriction enzyme used for cloning, encoded protein, forward or reverse orientation of the primer (F or R); QRT stands for qRT-PCR primers, PE stands for Primer Extension primers and RACE stands for Rapid Amplification of cDNA Ends primers) after the / symbol it is indicated the vector where the PCR product have been cloned.

| **No.** | **Brief description** | **Sequence (5ˊ-3ˊ)** | **PB code** |
| --- | --- | --- | --- |
| P1 | XbaI.P*tagB1.*F-225T | AAGGCCtctagaAGTTGGATGAAACCGGCTG | PB0033 |
| P2 | HindIII.P*tagB1.*R-225T | GGCCGGaagcttACGGTGGCAAGGTAATCC | PB0036 |
| P3 | BamHI.P*tac-tssE1.*F-225T | AAAAAAGGATCCttgacaattaatcatcggctcgtataatgGCGGCTGAAAGGCCGTTGTG | PB0722 |
| P4 | HindIII.P*tac-tssE1.*R-225T | TTTTTTaagcttATGCTGGCCTTGTACTCGGCC | PB0723 |
| P5 | XbaI.P*tac-tssJ1.*F-225T | AAAAAATCTAGAttgacaattaatcatcggctcgtataatgAACATCCTGTCCCAGACCC | PB0828 |
| P6 | HindIII.P*tac-tssJ1.*R-225T | GTGTGTaagcttTGATACACGCGCATCACC | PB0357 |
| P7 | XbaI.P*tac-hcp1.*F-225T | atatatTCTAGAttgacaattaatcatcggctcgtataatgGGACTACCTGTATGCCAGCC | PB0371 |
| P8 | HindIII.P*tac-hcp1.*R-225T | GATAGAaagcttGCCTTGTCTTCACCACTCTGG | PB0372 |
| P9 | XbaI.P*hcp4.*F-225T | TATATAtctagaCATGAAGTGCTTCATCGCCG | PB0690 |
| P10 | HindIII.P*hcp4.* R-225T | AGCTGTaagcttACTTCCTTGAGATGCGGGTC | PB0691 |
| P11 | XbaI.P*hcp5.*F-225T | TATAAAtctagaCATCGAAACTGACCAAATCGTCATC | PB0829 |
| P12 | HindIII.P*hcp5.*R-225T | TTTTTTaagcttAGCTTCACTTCCTTCAGGTGC | PB0831 |
| P13 | XbaI.gacS-UP.F | ATATATtctagaCAACTCGGCCATGCTGAATGG | PB0494 |
| P14 | gacS-UP.R | agcgctcaaATCGAGCACACTCGCCTCC | PB0495 |
| P15 | gacS-DOWN.F | gtgctcgatTTGAGCGCTTGAGGGGATCTG | PB0496 |
| P16 | BamHI.gacS-DOWN.R | ATATATggatccCCATGATGCGAGCGAAGGC | PB0497 |
| P17 | XbaI.tke7-V5-UP.F | AATTAAtctagaGGGCTGGTACTGCTAGTGAC | PB0714 |
| P18 | Tke7-V5-UP.R | aggcttacccgtagaatcgagaccgaggagagggttagggataggcttaccTCGCACTGCCTCATACAAAATCTC | PB0715 |
| P19 | Tke7-V5-DOWN.F | gattctacgggtaagcctatccctaaccctctcctcggtctcgattctacgtgaTATGAGGCAGTGCGATGAGTAAC | PB0716 |
| P20 | BamHI.tke7-V5-DOWN.R | TTGGTTggatccATCACACACACGCTGATTGC | PB0717 |
| P21 | KpnI.hcp5-strepII.F-234 | AGTTAAggtacctaacaggaggaattaaccATGACGATGTCTGAACATCGTCG | PB705 |
| P22 | XbaI.hcp5-strepII.R-234 | AGATCTtctagatcacttttcgaactgcgggtg | PB706 |
| PB23 | SEVAF | GGGTTTTCCCAGTCACGAC | PB0003 |
| PB24 | SEVAR2 | ACAATTTCACACCCTAGGCC | PB0004 |
| P25 | XbaI.hcp1.F-KT25 | caggaaacagctATGTTGTTAATGGAGAGTTTCACAATGGC | PB0394 |
| P26 | BamHI.hcp1.R-KT25 | AAAAGGATCCTTAGGCAAACACTTTGTTC | Not in our database |
| P27 | KT25.F | ATCTGTCCAACTTCCGCGAC | PB634 |
| P28 | KT25.R | TGTGCTGCAAGGCGATTAAG | PB635 |
| P29 | XbaI.hcp5.F-T18 | AATTCATCTAGAGacgatgtctgaacatcgtcg | PB0880 |
| P30 | BamHI.hcp5.R-T18 | TATTAAGGATCCTCcgcatacttcttgttgttgatgc | PB0881 |
| P31 | T18.F | ATGCTTCCGGCTCGTATGTTGTG | PB0091 |
| P32 | T18.R | TGCGGAACGGGCGCCGGCGCGAGCG | PB0090 |
| P33 | NheI.tke7.F-238DM | ATTAATGCTAGCgaggcgcgcagcttcattaatc | PB0832 |
| P34 | BamHI.tke7.R-238DM | TTTTTTGGATCCgtgcaagttactcatcgcactgcc | PB0833 |
| P35 | PelB-tke7.F | CTAGCatgaagtacctgctgccgaccgcggcggcgggtctgctgctgctggcggcgcagccggcgatggcgG | PB0877 |
| P36 | PelB-tke7.R | CTAGCcgccatcgccggctgcgccgccagcagcagcagacccgccgccgcggtcggcagcaggtacttcatG | PB0878 |
| P37 | NheI.tke2.F-238DM | ATTAATGCTAGCcgttatgtcactcaggaccctattgg | PB0879 |
| P38 | BamHI.tke2V5.R-238DM | TTTTTTGGATCCtcacgtagaatcgagaccgaggagag | PB0834 |

#### Table S4. Putative Rsm binding sites (located near ribosomal binding sites) identified *in silico* within the T6SS genomic regions of *P. putida* KT2440 (Dataset 1) and in the experimental assay described in (Huertas-Rosales *et al.*, 2021). In this study, KT2440 mRNA complexed with RsmA, RsmE and RsmI proteins was purified by affinity chromatography and sequenced to identify genes that could be regulated at the post-transcriptional level. The table compiles information on the loci conforming the K1-T6SS clusters and the *hcp4* and *hcp5* orphan genes. Consensus sequences: 5’ -A/UCANGGANGU/A -3’ 5′- RUACARGGAUGU-3′ (R=G/A)

| Locus name | Rsm binding site | | Sequence | Strand | coordinates | coordinates |
| --- | --- | --- | --- | --- | --- | --- |
| **Structural operon** | | | | | | |
| PP5562 | *tagB1* | GCACAUGGACGA | | reverse | 3497589 | 3497600 |
| PP3098 | *tssE1* | GUACAGGGAUGA | | reverse | 3494688 | 3494699 |
| PP3094 | *tssJ1* | GUACCCGGAGGU | | reverse | 3488739 | 3488750 |
| PP3089 | *hcp1.1** | GUUAAUGGAGAG | | reverse | 3478587 | 3478598 |
|  | *hcp1.2* | AGUACUGGAUGA | | reverse | 3478524 | 3478535 |
|  | *hcp1.3* | ACACAAGGACUG | | reverse | 3478509 | 3478520 |
|  | *hcp1.4* | CGACAUGGAAGA | | reverse | 3478266 | 3478277 |
| *hcp4* orphan | | | | | | |
| PP0655 | *hcp4.1* | UACCAAGGAGAG | | forward | 763003 | 763014 |
|  | *hcp4.2* | AGCGCUGGAUGA | | forward | 763065 | 763076 |
|  | *hcp4.3* | ACAUCGGGACGT | | forward | 763165 | 763176 |
|  | *hcp4.4* | GCGGCUGGAAGC | | forward | 763230 | 763241 |
| *hcp5* orphan | | | | | | |
| PP4886 | *hcp5.1* | ACCUCAGGAUGA | | reverse |  |  |
|  | *hcp5.2* | UACCAGGGAGAG | | reverse |  |  |
|  | *hcp5.3* | GCUGCUGGAGGC | | reverse |  |  |
|  | *hcp5.4* | CCUGAAGGAAGU | | reverse |  |  |
|  | *hcp5.5* | CGUCCUGGAAGA | | reverse |  |  |
|  | *hcp5.6* | GCCAGUGGAAGU | | reverse |  |  |
|  | *hcp5.7* | ACGGGCGGAUGG | | reverse |  |  |

### LEGENDS FOR SUPPLEMENTARY INFORMATION DATASETS

Dataset S1. **Putative Rsm binding sites within the T6SS genomic regions of *P. putida* KT2440.** Overview of the results originating from an *in silico* analysis of the putative Rsm binding sites located within the T6SS genomic regions of *P. putida* KT2440. The T6SS genomic regions are organised by clusters based on the description by (Bernal *et al.*, 2017) (clusters K1-T6SS, K2-T6SS, K3-T6SS and orphans *hcp4*, *hcp5*, *hcp6*, *vgrG4* and *vgrG5* are shown in different tabs). The search for Rsm binding sites was performed using the 12-bps domains NNNNNN**GGA**N**G**N / NNN**CA**N**GGA**NNN based on the most conserved bases of previously characterised consensus sequences 5’-A/U**CA**N**GGA**N**G**U/A-3’ and 5′-RUA**CA**R**GGA**U**G**U-3′. The putative Rsm binding sites identified in the same strand and direction of the T6SS genes are depicted. Min and Max referred to the coordinates of the sequence in the reference strain KT2440 (NC_002947). Rsm binding sites within a locus are labelled with the locus name, while the ones within intergenic regions are indicated as this. Colum J showed the binding targets of RsmIEA as identified in (Huertas-Rosales *et al.*, 2021) by RAP-Seq. This column is colour-coded according to the Rsm protein that is bound to that region. Pink cells indicate the RNA-binding target of RsmA, while blue and green cells indicate the RNA target binding of RsmE and RsmI, respectively. RNA targets shared by more than one Rsm protein are coloured with the fusion of the previous colours. Last tab (in blue) named Huertas-Rosales 2021 showed the subset of RAP-Seq data related to T6SS clusters extracted from the supplementary information of this reference. The tabs in red (K1-T6SS, *hcp4* and *hcp5*) highlighted the T6SS clusters where Rsm targets were identified by RAP-Seq.

### SUPPLEMENTARY REFERENCES

Annette C. Vergunst, Barbara Schrammeijer, Amke den Dulk-Ras, Clementine M. T. de Vlaam, Tonny J. G. Regensburg-TuÏnk, and Paul J. J. Hooykaas (2000) VirB/D4-Dependent Protein Translocation from *Agrobacterium* into Plant Cells. *Science (1979)* **290**: 979–982.

Bernal, P., Allsopp, L.P., Filloux, A., and Llamas, M.A. (2017) The *Pseudomonas putida* T6SS is a plant warden against phytopathogens. *ISME Journal* **11**: 972–987.

Bernal, P., Furniss, R.C.D., Fecht, S., Leung, R.C.Y., Spiga, L., Mavridou, D.A.I., and Filloux, A. (2021) A novel stabilization mechanism for the type VI secretion system sheath. *Proc Natl Acad Sci U S A* **118**: e2008500118.

Boyer, H.W. and Roulland-Dussoix, D. (1969) A complementation analysis of the restriction and modification of DNA in Escherichia coli. *J Mol Biol* **41**: 459–72.

Calles, B., Goñi‐Moreno, Á., and Lorenzo, V. (2019) Digitalizing heterologous gene expression in Gram‐negative bacteria with a portable ON/OFF module. *Mol Syst Biol* **15**: e8777.

Hanahan, D. (1985) Techniques for transformation of E. coli. In *DNA Cloning: A Practical Approach*. Glover, D.M. (ed). Virginia, p. 109.

Herrero, M., Lorenzo, V. de, and Timmis, K.N. (1990) Transposon vectors containing non-antibiotic resistance selection markers for cloning and stable chromosomal insertion of foreign genes in gram-negative bacteria. *J Bacteriol* **172**: 6557–6567.

Huertas-Rosales, Ó., Romero, M., Chan, K.-G., Hong, K.-W., Cámara, M., Heeb, S., et al. (2021) Genome-Wide Analysis of Targets for Post-Transcriptional Regulation by Rsm Proteins in *Pseudomonas putida*. *Front Mol Biosci* **8**: 624061.

Huertas-Rosales, O., Romero, M., Heeb, S., Espinosa-Urgel, M., Amara, M.C., and Ramos-Gonz Alez, I. (2017) The *Pseudomonas putida* CsrA/RsmA homologues negatively affect c-di-GMP pools and biofilm formation through the GGDEF/EAL response regulator CfcR. *Environ Microbiol* **19**: 3551–3566.

Kaniga, K., Delor, I., and Cornelis, G.R. (1991) A wide-host-range suicide vector for improving reverse genetics in Gram-negative bacteria: inactivation of the blaA gene of Yersinia enterocolitica. *Gene* **109**: 137–141.

Karimova, G., Pidoux, J., Ullmann, A., and Ladant, D. (1998) A bacterial two-hybrid system based on a reconstituted signal transduction pathway. *Proceedings of the National Academy of Sciences* **95**: 5752–5756.

Kessler, B., Lorenzo, V. de, and Timmis, K.N. (1992) A general system to integrate *lacZ* fusions into the chromosomes of gram-negative eubacteria: regulation of the *Pm* promoter of the TOL plasmid studied with all controlling elements in monocopy. *Mol Gen Genet* **233**: 293–301.

Martínez-García, E., Fraile, S., Algar, E., Aparicio, T., Velázquez, E., Calles, B., et al. (2023) SEVA 4.0: an update of the Standard European Vector Architecture database for advanced analysis and programming of bacterial phenotypes. *Nucleic Acids Res* **51**: D1558–D1567.

Nakazawa, T. (2002) Travels of a *Pseudomonas*, from Japan around the world. *Environ Microbiol* **4**: 782–786.
